# Social integration predicts survival in female white-faced capuchin monkeys

**DOI:** 10.1101/2020.08.04.235788

**Authors:** Kotrina Kajokaite, Andrew Whalen, Jeremy Koster, Susan Perry

**Author notes:** Corresponding author: Kotrina Kajokaite, Department of Human Behavior, Ecology and Culture, Max Planck Institute for Evolutionary Anthropology, Deutscher Platz 6 04103 Leipzig, Germany.

## Abstract

Across multiple species of social mammals, a growing number of studies have found that that individual sociality is associated with survival. In long-lived species, like primates, lifespan is one of the main components of fitness. We used 18 years of data from the Lomas Barbudal Monkey project to quantify social integration in 11 capuchin (*Cebus capucinus*) groups and tested whether female survivorship was associated with females’ tendencies to interact with three types of partners: (1) all group members, (2) adult females, (3) adult males. We found strong evidence that females who engaged more with other females in affiliative interactions and foraged in close proximity experienced increased survivorship. We found some weak evidence that females might also benefit from engaging in more support in agonistic contexts with other females. These benefits were evident in models that account for the females’ rank and group size. Female interactions with all group members also increased survival, but the estimates of the effects were more uncertain. In interactions with adult males, only females who provided more grooming to males survived longer. The results presented here suggest that social integration may result in survival-related benefits. Females might enjoy these benefits through exchanging grooming for other currencies, such as coalitionary support or tolerance.

## INTRODUCTION

A key question for understanding the evolution of animal sociality is: Do more social individuals have higher fitness? It has been hypothesized that social animals have evolved cognition characterized by skills and motivations to monitor their social world and interact with conspecifics in fitness-enhancing ways (Jolly, 1966; Humphrey, 1976; Whiten and Byrne, 1997). A number of studies link differences in the social behavior of individuals to components of reproductive success, such as fertility (Fedigan et al., 2008; Schülke et al., 2010; Gilby et al., 2013; Feldblum et a. 2021) and offspring survival (Silk, 2009; Silk et al., 2003; Kalbitzer et al., 2017). In long-lived, iteroparous species, lifespan is also an important component of life-time reproductive success that contributes to variation in fitness (Clutton-Brock, 1998), and there is evidence that aspects of individual sociality are associated with lifespan in humans and nonhuman animals (Snyder-Mackler et al., 2020).

Much research has investigated whether social relationships that primates develop with some social partners predict longevity (reviewed in Snyder-Mackler et al. 2020). Strong and durable social bonds with either female (Silk et al. 2010; Archie et al. 2014) or male (Archie et al. 2014) partners predict adult female longevity. Some evidence suggests that the number of weak bonds is also predictive of fitness-related benefits (McFarland et al. 2017; Silk et al. 2018). Studying social bonds helps us identify the functional aspects of cultivating social relationships with specific partners (Snyder-Mackler et al. 2020), but this approach might not account for important interindividual variation in affiliation tendency, number of partners, and number of strong bonds (Ostner and Schülke, 2018).

Here we focus on an individual’s social integration in their group. In human studies these measures are called “structural”, with the emphasis being on the individual’s position in the social network (Snyder-Mackler et al. 2020). Survival-related benefits flow to socially integrated individuals through increased social tolerance, better access to resources, and/or beneficial spatial location that results in reduced predation or minimizes the risk of injury during coalitionary conflicts (Ellis et al. 2019). Various versions of social integration measures have been shown to predict longevity in some species (e.g., feral horses, *Equus caballus*: Nuñez et al. 2015; rhesus macaques, *Macaca mulatta:* Brent et al. 2017; chacma baboons, *Papio ursinus:* McFarland et al. 2017).

In this paper, we ask how an adult white-faced capuchin (*Cebus capucinus*) female’s survival is predicted by her social integration into her group. Capuchin monkeys live in stable, well-defined groups (Perry and Manson, 2009; Perry, 2012). Capuchin females, being the philopatric sex, have an opportunity to develop strong and enduring social relationships with other females with whom they co-reside for their entire lives. Male capuchins migrate multiple times, and their tenure in the same social group can vary from approximately 2 weeks to 18 years, making it hard to distinguish between death and dispersal outside the study area (Perry 2012). In contrast, the longevity and lifetime reproductive success of adult females can be accurately documented, and are tightly associated. Female capuchins give first birth when they are 6 years and then have 2-year interbirth intervals (Perry 2012). In this population, eighteen females who lived more than 20 years (maximum 37 years), had an average lifetime reproductive success of 8 offspring (SD = 2.12, range = 5 – 13; Perry 2012). Adult males outrank all adult females and are physically larger, which makes them valuable coalitionary partners (Perry and Manson 2008). To account for these differences in opportunities to form relationships, we estimate an adult female’s social integration with other adult females and with adult males separately. In addition, we estimate adult females’ social integration using all observed interactions that they had with all of their group members (adult females, adult males and immatures) in order to estimate as accurately as possible the effects of their overall social integration within their social group on survival.

To do this, we chose an individual-level metric, by quantifying the adult female’s base rate of engaging in a behavior with three types of partners: (1) all group members, (2) adult females, (3) adult males. We assume that participating more in some social interactions translates into more survival-related benefits, such as increased access to resources, reduced predation and/or wounding risk, and increased rank (Ostner & Schülke 2018; Thompson 2019; Ellis et al. 2019). The pathways from the interactions to survival-related benefits can be either direct or manifested through exchange of beneficial social interactions with others (Ostner & Schülke 2018; Thompson 2019).

We aimed to quantify social integration in different contexts of capuchin social life: (1) affiliative interactions (measured via grooming), (2) support in agonistic contexts, and (3) one form of tolerance in feeding context (measured by time spent foraging in proximity, although this does not assume that foraging near others is a proxy for tolerance from group members in non-foraging contexts). We quantified interactions in each domain separately and, where possible (i.e., for grooming and coalitionary support), we estimated a female’s role as both a provider and as a receiver of the behavior. If females are trading some services for others, their role as a receiver and a giver might be associated with a different pathway to survival-related benefits (*see supplemental material for detailed descriptions of possible pathways*).

In this paper, we examine whether females’ participation in affiliative interactions, support in agonistic contexts, or spatial association during feeding (analyzed separately for (a) all group members, (b) same-sex, and (c) opposite-sex) is a predictor of survival in wild white-faced capuchin monkeys. We used demographic and behavioral data spanning 18 years, from the Lomas Barbudal Monkey Project dataset. We employed a time-varying statistical approach to take into account how social integration changes through time and tested whether it predicts survival in 132 adult females.

We hypothesize that females who are more socially integrated experience survival-related benefits. Specifically, we predicted that the more females provide and receive either grooming or coalitionary support, the better they will survive. We also predicted that females who are more likely to forage in close proximity of others will survive longer. We expect to find support for these predictions across all partner types (all group members, adult females, adult males), but we anticipate that social integration with adult female partners will most strongly predict survival.

## METHODS

### Study subjects and the datasets

We studied members of the wild white-faced capuchin population at the Lomas Barbudal Biological Reserve and surrounding private lands in Guanacaste, Costa Rica (Perry et al., 2012). The dataset has longitudinal records including demographic information, pedigree information, and social interactions on individuals living in 11 capuchin social groups. The data on capuchin behavior were collected between January 2002 and December 2019. All groups in this study were observed during at least 7 calendar years (mean = 13.3, SD = 4.4, range = 7 - 18). The behavioral and demographic data on each group were collected by experienced observers during visits lasting at least 6 hours/day. Supplemental materials provide more demographic information about each group (*Section 5, Table S2*).

The primary subjects of this analysis were 132 adult females (adulthood defined as beginning at 5 years of age). Sixty-two females died during the study period, and we have information about the causes of death in 10 of those cases (*see supplemental materials, Section 2 Table S1, for causes of death and distribution across age categories*).

### Measuring Social Integration

We treated grooming and coalition formation as directed behaviors, and we used observations of individuals as both initiators and recipients of the behavior. We did not have information about which individual had initiated the proximity when foraging, and therefore foraging in proximity was treated as an undirected behavior. We treated each of the five interaction types (*grooming giving, grooming receiving, support giving, support receiving, foraging*) as a separate measure of social integration.

In calculating the frequency with which adult females engaged in these interactions, we assembled three datasets. The first dataset included adult female interactions with all group members (adult males, adult females and immatures). The second dataset included adult female interactions with other adult females only, and the third dataset included adult female interactions with adult males only. These latter datasets exhibit heterogeneous samples sizes because of imbalanced sampling of groups and individuals throughout the study (Table 1). We estimated the posterior mean and standard deviation of each individual-level social integration measure and propagated that measurement uncertainty when estimating the effects of sociality on survival (*see supplemental material, Section 3*).

**Table 1.** Summary statistics of the three datasets. Each dataset contains social integration measures estimated from interactions between adult females and their social partners: (1) all group members, (2) adult females only, (3) adult males only.

| Dataset | # females | # female years | # deaths |
| --- | --- | --- | --- |
| All group members | 132 | 1058 | 62 |
| Adult females | 132 | 985 | 55 |
| Adult males | 129 | 1002 | 55 |

### Grooming (grooming giving and grooming receiving)

Grooming rates were estimated using data collected during 10-minute focal follows. To estimate individual grooming rates, we calculated dyadic counts of grooming and dyadic opportunities for grooming. The opportunity for an *ij* dyad to engage in grooming was calculated as the sum of the focal follows of *i* and the focal follows of *j* at times when *i* and *j* were co-resident. A count of 1 was assigned if *i* groomed *j* at least once during a focal follow; otherwise 0 was assigned. The same was done when evaluating if *j* groomed *i*. There were 116,537 focal follows in our dataset (an average of 35 per individual per year and an average of 782 per group per year). We observed a total of 38,862 dyadic grooming events, with a median of 2 grooming giving events per individual per year and much individual variation (range = 0 – 288, where in 1231 instances, individuals were not observed providing grooming in a particular year), and with a median of 4 grooming receiving events (range = 0 – 143, where in 816 instances, individuals were not observed receiving grooming in a particular year).

### Joining an ongoing conflict (support giving and support receiving)

The behavior of joining a coalitionary conflict was defined as an individual intervening on one side during an ongoing aggressive conflict. This definition indicates only the functional aspect of joining a side; it entails no inferences about internal psychological states such as the intent to help a specific individual. Since aggressive interactions are salient and harder to miss than quiet activities like grooming, aggressive interactions were collected both *ad libitum* and during focal follows. The types of aggressive signals that qualified as support included physical aggression directed at an opponent, chases, signals of aggressive intent (including facial expressions, vocalizations and gestures) and postures and gestures of coalitionary support with a partner vs. the partner’s opponent (e.g., overlords and cheek to cheek postures). The chronological stream of aggressive behaviors was divided into 5-minute intervals. In order to identify instances of joining a coalitionary conflict, monkey *i* is identified as joining monkey *j* if *i* performed an aggressive behavior toward either monkey *j*’s opponent or victim within the context of the intervals. The measure is dichotomous, and a single instance was recorded for the occasions when there were multiple observations of monkey *i* joining monkey *j* during the interval. To calculate the opportunities to join a coalitionary conflict, all individuals who were co-resident during the aggressive conflict were regarded as having the opportunity to join on either side during the conflict. There were 48,081 5-minute aggressive intervals in our dataset (an average of 14 per individual per year and an average of 322 per group per year). We observed a total of 38,862 dyadic support events, with a median of 4 support giving events per individual per year and much individual variation (range = 0 – 241, where in 939 instances, individuals were not observed providing grooming in a particular year), and with a median of 4 support receiving events (range = 0 – 239, where in 893 instances, individuals were not observed receiving grooming in a particular year).

### Foraging

*Foraging* was estimated from group scans that occurred in the context of foraging. In group scans, the identity of the scanned individuals, their activity and their proximity to other individuals within 10 body lengths (∼2 m) was noted. We considered individuals to be foraging in close proximity if they were scanned within 5 body lengths (∼1m) of each other. For each dyad, we scored whether they were observed foraging within close proximity in 10-minute intervals. The number of opportunities that the dyad had to forage within close proximity is a sum of group scans in the foraging context that are 10 minutes apart, where one of the individuals is a subject of a group scan. There were 310,910 group scans in our dataset (an average of 93 per individual per year and an average of 2086 per group per year). We observed a total of 100,532 cases of dyadic foraging in close proximity, a median of 16 per individual per year and much individual variation (range = 0 – 217, where in 248 instances individuals were not observed foraging in close proximity of others in a particular year).

### Individual social integration measures

The data for these analyses were collected across eighteen years and the number of observed social groups and individuals generally increased over time. As a result, the density of data is uneven across time periods, social groups, and individuals. We incorporated uneven distributions of the data by aggregating the data annually and using adaptations of the multilevel Social Relations Model (Snijders and Kenny, 1999; Koster et al. 2020) to estimate individual annual estimates of grooming, coalitionary support, and foraging in close proximity (*see supplemental material, Section 3*). This method provided estimates of individual social integration that reflect measurement uncertainty, with the uncertainty increasing for infrequently observed individuals.

Social integration estimates from the Social Relations Model can be conceptualized as a female’s base rate of either providing or receiving behavior, after accounting for average rates for the population. It describes how likely a female is to participate in an interaction (either as a recipient or as a provider, if applicable) with another individual in her social group in a given year. The measure is estimated as a random effect in the statistical model and centered on zero, which conceptually represents an average monkey in the population. The social integration estimate then is an offset from the population average (population here depends on the dataset: (1) all individuals, (2) only adult females, (3) only adult females and males). In general, our social integration measures are similar to nodal in- and out-degree centrality in the social networks tradition (Hasenjager and Dugatkin, 2015).

### Modeling survival as a function of the individual social integration measure

To investigate whether sociality is associated with adult female survival, we used Bayesian accelerated failure time models (Wei 1992; *see supplemental materials, Section 7*). In separate models, each of the five annual individual social integration measures (mean and SD) was modeled as a predictor of survival probability over one-year periods. In the main text, we report models in which survival in year *t* is modeled as a function of social integration estimated in the same year. As a supplemental analysis, we model survival in year *t* as a function of social integration in the preceding year, *t-1*. The former approach has a larger sample size whereas the second approach assumes that the benefits of social integration are evident over an intermediate timeframe (*see supplemental materials, Section 10*).

Survival rate is age-dependent in capuchin monkeys (Colchero et al. 2021). Among adults, survival decreases with age and the rate appears to be approximately linear in our population (Colchero et al. 2021; *supplemental materials, Section 2, Table S1*). We included an age effect in our model to account for this, and it allowed us to focus on the main covariates of interest in these analyses – social integration measures – and test if they predict adult female survival. We use a time-varying statistical approach to account for age-related changes in individual social integration (*see supplemental materials, Section 6)* and in the availability of social partners (Archie et al. 2014). These models included the following time-varying (calendar year-specific) covariates: female’s dominance rank (ranges from 0 to 1, where 1 represents the highest rank; *see supplemental material, Section 5*), the average number of individuals in her group (including juveniles and infants) throughout the year (*see supplemental material Section 5 for further details on covariates*). There are alternative ways to model age dependent survival, such as using a Weibull model (Carroll 2003), but the accelerated failure time model used here allows us to capture the effect of age (as a linear covariate) and consider time dependent changes in other covariates. To account for imbalanced sampling of groups and individuals, we also estimated unique random intercepts for each individual and for each group. In summary, we ask whether females who are of the same age, rank and group size, but differ in social integration measures have a different probability of survival. i.e., Are those with greater social integration measures less likely to die that year?

We used a Bayesian approach to fit the accelerated failure time model (Wei 1992). All of the covariates were standardized by subtracting the mean and dividing it by the standard deviation. Models were run using Stan (v.2.19.1) and the *rethinking* package (v. 1.93: McElreath, 2020) in *R* (v. 3.6.2; R Core Team 2019).

## RESULTS

### Individual social integration measures

For perspective on the individual social integration measures, we present female posterior mean estimates’ minima and maxima in Table 2, which shows the range for individual social integration measures by domains: (1) *grooming giving*, (2) *grooming receiving*, (3) *support giving*, (4) *support receiving*, (5) *foraging;* and by the dataset: (a) all group members, (b) adult females, (c) adult males. Females varied the most in how much grooming they provided (range: -3.17; 3.45) and varied the least in how much grooming they received when grooming exchanged among all group members were considered (range: -0.99; 0.87; see Table 2).

**Table 2.** The ranges of individual social integrations measures from nine Social Relation Models. The estimates represent three Social Relations Models which estimated five individua social integration measures: (1) *grooming giving*, (2) *grooming receiving*, (3) *support giving*, (4) *support receiving*, (5) f*oraging*, for each dataset: (a) all group members, (2) adult females, (3) adult males. The reported quantities are female posterior minima and maxima.

| Social integration measure | All group members | Adult females | Adult males |
| --- | --- | --- | --- |
| Grooming giving | -3.17; 3.45 | -2.42; 1.63 | -1.59; 2.67 |
| Grooming receiving | -0.99; 0.87 | -1.63; 0.88 | -1.22; 1.56 |
| Support giving | -2.10; 2.86 | -1.95; 1.94 | -2.10; 1.90 |
| Support receiving | -1.15; 1.43 | -0.95; 0.94 | -1.13; 1.06 |
| Foraging | -2.59; 1.97 | -1.71; 1.94 | -1.82; 1.94 |

### Affiliative interactions and survival

Social integration measured via grooming reliably predicted survival when the measure was females’ grooming with adult females (*grooming giving:* ß = 0.30, CI: 0.07, 0.54; and *grooming receivin*g: ß = 0.29, CI: 0.07, 0.51; see Table 3). When we measured female social integration into her social group overall (grooming interactions with adults and immatures), *grooming giving* predicted survival reliably (ß = 0.54, CI: 0.35, 0.73; see Table 3), and *grooming receiving* showed marginal support (ß = 0.21, CI: 0.00, 0.41; see Table 3) since the credible interval of the coefficient is mostly above zero. We also found marginal support that female grooming interactions with adult males predicted survival when females provided grooming (*grooming giving:* ß = 0.20, CI: -0.04, 0.45; see Table 3). We found no evidence that females who received grooming from adult males survived better. The effect of *grooming receiving* from adult males was negative (ß = –0.10, CI: -0.31, 0.10; see Table 3), but the effect size is small and the differences in probability of dying are small for those who receive a lot of grooming from adult males in comparison to those who receive little grooming from adult males (see Figure 1). Figures 1 illustrates the accelerated failure time model predictions of probability of dying in a given year (*see supplemental materials, Section 8, for more details on the model results*).

**Table 3.** Fifteen accelerated failure time models predicting female survival as a function of female’s social integration via (1) *grooming giving*, (2) *grooming receiving*, (3) *support giving*, (4) *support receiving*, and (5) *foraging* to (a) all group members (N=132 females, 1058 female-years of data, 66 censored cases), to (b) adult females (N=132 females, 985 female-years of data, 77 censored cases), and to (c) adult males (N=129 females, 1002 female-years of data, 74 censored cases). The reported quantities are posterior means (89% credible intervals in parentheses).

| Model | All group members | Adult females | Adult males |
| --- | --- | --- | --- |
| **(1) Grooming giving** |  |  |  |
| Intercept | 9.46 (8.91; 10.02) | 9.41 (8.88; 9.96) | 9.46 (8.92; 10.01) |
| Grooming giving | 0.54 (0.35; 0.73) | 0.30 (0.07; 0.54) | 0.20 (-0.04; 0.45) |
| Age | -0.07 (-0.10; -0.04) | -0.07 (-0.1; -0.04) | -0.08 (-0.11; -0.05) |
| Rank | -0.24 (-0.50; 0.01) | -0.14 (-0.41; 0.13) | -0.05 (-0.31; 0.21) |
| Group size | 0.15 (-0.11; 0.42) | 0.09 (-0.18; 0.38) | 0.10 (-0.17; 0.37) |
| **(2) Grooming receiving** |  |  |  |
| Intercept | 9.48 (8.95; 10.01) | 9.43 (8.89; 9.99) | 9.52 (8.96; 10.06) |
| Grooming receiving | 0.21 (0.00; 0.41) | 0.29 (0.07; 0.51) | -0.10 (-0.31; 0.10) |
| Age | -0.09 (-0.12; -0.06) | -0.07 (-0.10; -0.05) | -0.09 (-0.12; -0.06) |
| Rank | -0.13 (-0.39; 0.12) | -0.15 (-0.42; 0.12) | 0.02 (-0.24; 0.28) |
| Group size | 0.12 (-0.15; 0.38) | 0.09 (-0.18; 0.38) | 0.07 (-0.21; 0.34) |
| **(3) Support giving** |  |  |  |
| Intercept | 9.50 (8.97; 10.05) | 9.46 (8.94; 10.02) | 9.52 (8.98; 10.07) |
| Support giving | 0.15 (-0.12; 0.42) | 0.25 (-0.03; 0.53) | -0.14 (-0.40; 0.12) |
| Age | -0.09 (-0.12; -0.06) | -0.08 (-0.11; -0.05) | -0.09 (-0.12; -0.06) |
| Rank | -0.16 (-0.46; 0.13) | -0.19 (-0.48; 0.11) | 0.07 (-0.20; 0.35) |
| Group size | 0.12 (-0.14; 0.38) | 0.08 (-0.20; 0.36) | 0.05 (-0.23; 0.34) |
| **(4) Support receiving** |  |  |  |
| Intercept | 9.49 (8.96; 10.03) | 9.45 (8.89; 9.99) | 9.53 (8.98; 10.07) |
| Support receiving | 0.21 (-0.05; 0.45) | 0.23 (-0.04; 0.51) | -0.12 (-0.38; 0.12) |
| Age | -0.09 (-0.12; -0.06) | -0.08 (-0.11; -0.05) | -0.09 (-0.12; -0.06) |
| Rank | -0.20 (-0.49; 0.1) | -0.18 (-0.47; 0.13) | 0.07 (-0.21; 0.34) |
| Group size | 0.12 (-0.14; 0.38) | 0.08 (-0.19; 0.35) | 0.06 (-0.22; 0.35) |
| **(5) Foraging** |  |  |  |
| Intercept | 9.47 (8.92; 10.02) | 9.48 (8.94; 10.01) | 9.51 (8.95; 10.07) |
| Foraging | 0.28 (-0.01; 0.58) | 0.32 (0.06; 0.58) | 0.08 (-0.17; 0.34) |
| Age | -0.09 (-0.12; -0.06) | -0.08 (-0.11; -0.05) | -0.09 (-0.11; -0.06) |
| Rank | -0.18 (-0.45; 0.09) | -0.15 (-0.43; 0.12) | -0.02 (-0.3; 0.25) |
| Group size | 0.22 (-0.07; 0.51) | 0.13 (-0.16; 0.42) | 0.10 (-0.17; 0.38) |

\* See supplemental materials for more details on models' results.

**Figure 1.**
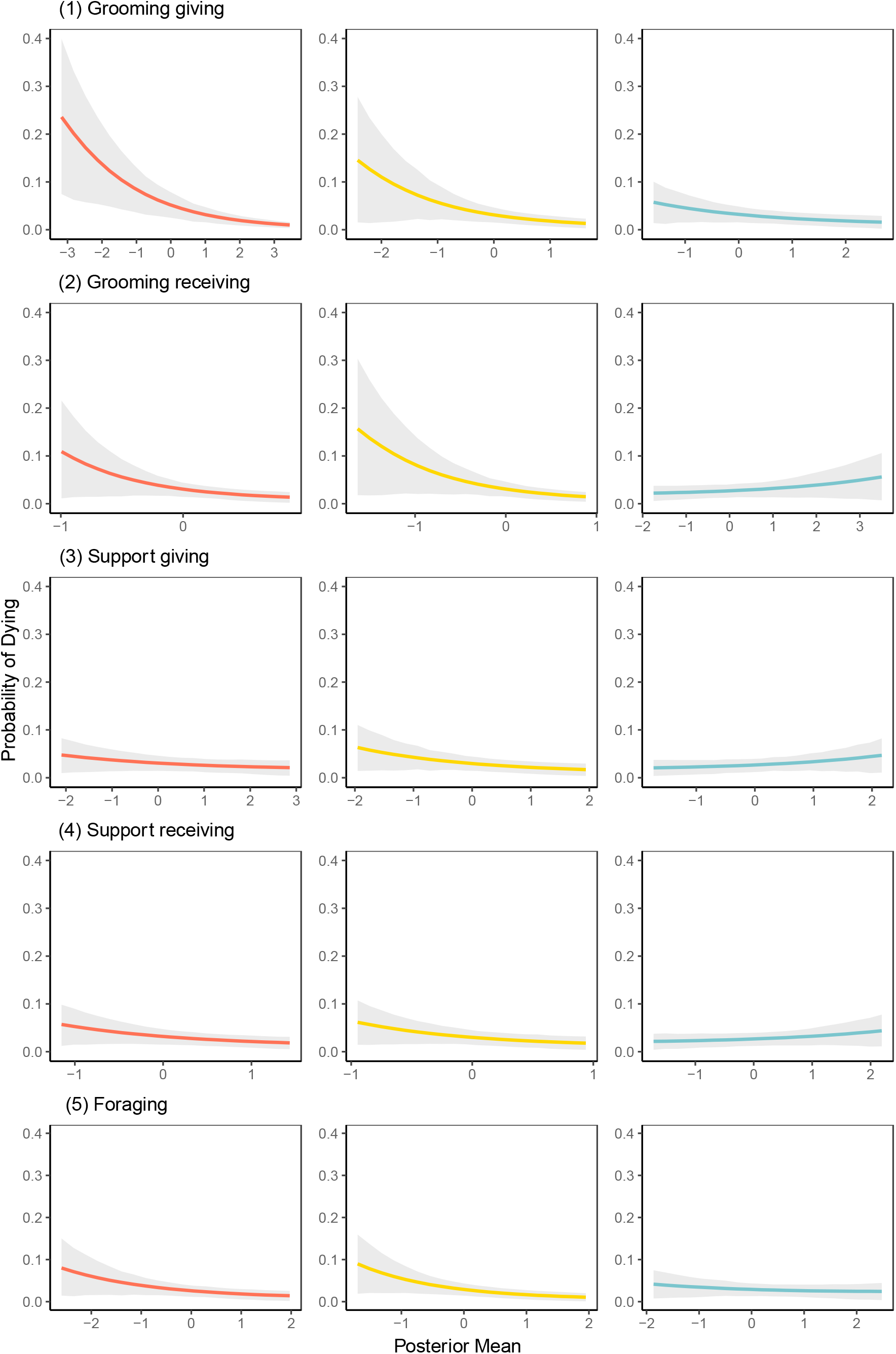
Fifteen accelerated failure time models’ predicted annual probabilities of dying as a function of the social integration measured through female’s (1) *grooming giving*, (2) *grooming receiving*, (3) *support giving*, (4) *support receiving*, and (5) *foraging* to: (a) all group members (red), (b) adult females (yellow), (c) adult males (blue). All other covariates are set to a sample mean. Grey region represents 89% credible interval. Note the different scales of the x-axes that reflect the differences in variance in individual social integration estimates.

### Support in agonistic contexts and survival

In comparison to affiliative interactions, we found weaker evidence that either providing coalitionary support or being a recipient of coalitionary support predicts survival. We found marginal support for increased survival being associated with (a) *support giving* to adult females (ß = 0.25, CI: -0.03, 0.53; see Table 3), (b) *support receiving* from adult females (ß = 0.23, CI: -0.04, 0.51; see Table 3), and (c) *support receiving* from anyone in one’s social group (ß = 0.21, CI: -0.05, 0.45; see Table 3). The credible intervals of these effects are mostly above zero. We found no evidence that support in agonistic interactions with adult males predicted survival among females.

### Foraging tolerance and survival

In foraging tolerance interactions, we found evidence that females who are more often observed foraging in close proximity to other adult females survive longer. The effect was reliably above zero (ß = 0.32, CI: 0.06, 0.58; see Table 3). We also found marginal support that females who forage in close proximity to anyone from their social group also survive longer (ß = 0.28, CI: -0.01, 0.58; see Table 3) since the credible interval of the effect is mostly above zero. Foraging near adult males had a positive posterior mean (ß = 0.08, CI: -0.17, 0.34; see Table 3), but the zero was comfortably within the credible interval suggesting that we have no strong evidence of these interactions predicting survival.

### Age and survival

To account for decreasing survival with age (Colchero et al. 2021; *supplemental materials, Section 2*), we included an age covariate in our models. As expected, in all of our models the age effect is reliably negative, indicating that it approximates decreasing survival as females age.

### Previous year’s social integration and survival

We found that previous year’s social integration estimates were less associated with survival in comparison to the models where survival in year *t* is modeled as a function of social integration estimated in the same year. Across all models and all datasets, the social integration effect sizes were smaller and more uncertain (*supplemental materials, Section 10*). Only the effect of *grooming giving* when measured through social integration with all group members exhibited a positive effect that was reliably above zero. This suggests that the benefits of social integration are more evident on a more immediate time frame.

## DISCUSSION

Female white-faced capuchin social behavior is associated with survivorship: social integration into affiliative interactions and feeding tolerance predicted adult female survival. We found marginal support that providing and receiving support in agonistic contexts is associated with survival, but the evidence was uncertain. Our results are broadly consistent with previous findings that variation in social behavior is associated with survival (Holt-Lunstad et al., 2010, Fagen and Fagen, 2004; Yee et al., 2008; Silk et al., 2010; Archie et al., 2014; Brent et al., 2017).

In our sample, five females appear to have died of malnutrition due to poor access to food during a drought, one was killed by a poacher, one disappeared when she was injured, and one vanished when she was pregnant. Two females died when hit by a car. For other deaths in our sample, we lack information about the causes, because most of the deaths are not observed (*see supplemental materials section 2 for more information on causes of death)*. In general, socially integrated females can experience reduced risk of predation or wounding from conspecifics, improved access to food resources, and improved social status (Ellis et al. 2019). From documented causes of death in this population, we have indirect evidence that access to these benefits can be beneficial to survival.

We considered female interactions with three types of partners: (1) all group members, (2) adult females, (3) adult males. We found the strongest evidence that social integration into female networks predicts survival. Females who frequently engaged in grooming and foraged in close proximity with other females experienced improved survivorship. We also found marginal support that females who frequently provided and received coalitionary support from other females possibly experienced survival-related benefits. The fact that female interactions with other females across all behavioral domains show at least some evidence of beneficial impact on survival suggests that females cultivate strong relationships with each other. This is consistent with findings in other female philopatric species: baboons (Silk et al. 2010; Archie et al. 2014) and macaques (Brent et al. 2017).

When we considered female interactions with male partners, we found virtually no evidence that heterosexual relationships are beneficial for female survival. The exception was a tendency for females who groom males to survive better, but this effect was uncertain. In grooming interactions between females and males, females are much more avid groomers and males rarely groom their partners and rarely reciprocate immediately (Perry 1997, 2012). Our finding in the grooming giving domain is consistent with the Archie et al. (2014) findings that adult female social connectedness via grooming of adult males (providing and receiving, in this case) predicted survival in wild female baboons. The lack of evidence that females experienced almost no survival-related benefits from their interactions with adult males might be due to males being the dispersing sex. The majority of the males have short tenures in social groups (Perry 2012), and as a result, females might not have enough time to develop relationships with them. However, females might benefit from strong relationships with a few male partners who co-reside with them for longer periods of time. For example, the alpha male has a special status in capuchin social groups and is often sought as a grooming and coalitionary partner by females and other group members (Perry 2008). Strong relationships with a long-term alpha might be beneficial to female survival.

We also considered a female’s overall integration into her social group via her interactions with all of her group members (adults and immatures); these results were similar to the survival-related effects when examining only interactions with other females. However, the effects on survivorship in this case were weaker and uncertain in receiving grooming, receiving coalitionary support, and foraging in close proximity. There was no evidence that providing coalitionary support to members of all age-sex classes is beneficial. Overall, this further provides evidence that female-female relationships have the greatest effect on female survivorship, although being more social in general might also provide some benefits.

What are the potential causal pathways connecting social behavior to survival-related benefits? Our study does not directly test any specific pathway of how social interactions contribute to survival. It is possible that females are pursuing multiple strategies, thereby clouding our efforts to see a single clear social strategy promoting survival. Social interactions might be directly advantageous: for example, feeding tolerance can provide better access to food. Or the survival-related benefits might accrue indirectly. For example, grooming might increase spatial centrality within the group, and as a result, centrally located females might avoid predation (Josephs et al., 2016). Or females might be exchanging interactions for other currencies (Noë and Hammerstein 1995), thereby gaining access to food (de Waal 1989, 1997; Tiddi et al., 2011; Jaeggi & Gurven 2013). We did find at least some evidence that female interactions with other females in all domains positively predicted survival, which suggests that females might be experiencing both direct and indirect benefits when interacting with each other.

Out of all behavioral domains we considered, we found the least support that participating in coalitionary conflicts results in improved survival. Coalitionary support can be either traded for itself or for other currencies (e.g., tolerance) which can increase female’s rank (Strauss and Holekamp 2019) and/or provide better access to food (Haunhorst et al. 2017). We found marginal support for females getting survival-related benefits, either direct or indirect, when providing and receiving support from other females. Joining a fight is potentially costly (e.g., risk of injury) and if there is reciprocity in coalitionary aggression (Kajokaite 2019), then frequent recipients might also experience costs associated with returning coalitionary support to their partners.

A possibility that our study does not address is that females might not only pursue multiple strategies, but also that these strategy sets might differ among the individuals. Some females might frequently engage in social interactions across all domains, interact with a wide range of partners, and trade some currencies for others. Some females might invest heavily in a few social partners and/or focus their efforts in some social currencies more than others. As a result, females might experience differential pay-offs based on their demographic and ecological conditions (McFarland et al. 2017).

Our analysis does not preclude possible confounders that could influence both social behavior and survival, in which case the observed correlations could be caused by other factors (e.g., a favorable genotype that promotes social behavior and survival) (Ostner and Schülke 2018). Also, although our supplemental models examine the effects of social integration and survival in successive years, it is possible that physical senescence induces a correlation between social behavior and survival on longer timescales. Our data suggest that social integration measures do slightly decrease with age; however, most females die before they reach 30 years of age, and therefore only a few of them are candidates for experiencing physical senescence-induced changes in social behavior (*see supplemental materials, Sections 2 and 6*).

## CONCLUSION

Across taxa, research on sociality has revealed diverse effects on components of fitness. This study examines sociality in detail, showing that affiliative interaction, feeding tolerance, and perhaps even support in agonistic contexts predict survivorship of adult female capuchins.

Therefore, this study adds a neotropical primate species to the list of mammalian species where a similar association between social integration and survival has been demonstrated. These results accentuate the need for greater attention to the mechanistic pathways that connect sociality to fitness (Ostner and Schülke, 2018; Thompson, 2019). As longitudinal data become increasingly common in studies of animal behavior, careful analyses can better elucidate the evolutionary consequences of variation in sociality among individuals.

## ETHICS

The study was strictly observational, and all protocols were approved by UCLA’s Animal Care Committee (protocol 2016-022). All necessary permits were obtained from SINAC and MINAE (the Costa Rican government bodies responsible for research on wildlife) and renewed every 6 months over the course of the study; the most recent scientific passport number being #117-2019-ACAT and the most recent permit being Resolución # M-P-SINAC-PNI-ACAT-072-2019. This research is in compliance with the Animal Behavior Society’s Guidelines for the Use of Animals in Research.

## DATA AND CODE ACCESSIBILITY

The data and code used to produce the analyses in this paper are available at: www.doi.org/10.5281/zenodo.5885634

## ACKNOWLEDGEMENTS

The following field assistants contributed significant amounts of data to the Lomas Barbudal Monkey Project dataset: C. Angyal, A. Autor, B. Barrett, L. Beaudrot, M. Bergstrom, R. Berl, A. Bjorkman, L. Blankenship, T. Borcuch, J. Broesch, D. Bush, J. Butler, F. Campos, C. Carlson, A. Cobden, M. Corrales, J. Damm, B. Davis, C. deRango, N. Donati, C. Dillis, G. Dower, R. Dower, A. Duchesneau, K. Feilen, J. Fenton, S. Fiello, K. Fisher, A. Fuentes J., M. Fuentes A., T. Fuentes, C. Gault, H. Gilkenson, I. Godoy, I. Gottlieb, J. Griciute, L.M. Guevara R., L. Hack, R. Hammond, S. Herbert, C. Hirsch, M. Hoffman, A. Hofner, C. Holman, J. Hubbard, S. Hyde, M. Jackson, O. Jacobson, E. Johnson, L. Johnson, S. Lee, S. Leinwand, T. Lord, M. Kay, E. Kennedy, D. Kerhoas-Essens, E. Johnson, S. Kessler, B. Krimmel, W. Lammers, S. Lopez, S. MacCarter, J. Mackenzie, M. Mayer, F. McKibben, W. Meno, A. Mensing, M. Milstein, C. Mitchell, Y. Namba, A. Neyer, C. O’Connell, J. C. Ordoñez, J., N. Parker, B. Pav, R. Popa, K. Potter, K. Ratliff, K. Reinhardt, N. Roberts Buceta, E. Rothwell, J. Rottman, H. Ruffler, S. Sanford, M. Saul, E. Seabright, I. Schamberg, S. Schembari, N. Schleissman, C. Schmitt, S. Schulze, A. Scott, S. Sita, J. Shih, K. Stewart, W. Tucker, E. Urquhart, K. van Atta, J. Vandermeer, L. van Zuidam, J. Verge, V. Vonau, A. Walker-Bolton, E. Wikberg, E. Williams, L. Wolf, and D. Wood. We thank H. Gilkenson and W. Lammers for long-term management of the field site during 2002-2013. We thank the Costa Rican park service (Sistema Nacional de Áreas de Conservación and Área de Conservación Tempisque), Hacienda Pelon de la Bajura, Hacienda Brin D’Amor, and the residents of San Ramon de Bagaces for permission to work on their land. D. Cohen assisted with database management and queries. This paper has benefited from helpful discussions with H. Clark Barrett, Donald Cohen, Joseph Manson, Mary Brooke McElreath, Oliver Schülke, and Julia Ostner. We thank three anonymous reviewers whose comments/suggestions helped improve and clarify this manuscript.

