## Supplemental materials for "Social integration predicts survival in female white-faced capuchin monkeys"

### Supplemental material

KAJOKAITE ET AL.

This supplemental document contains ten sections. First, we discuss possible pathways connecting each interaction type and role to a different benefit associated with survival. Second, we report what we know about causes of deaths in capuchin females. Third, we provide a description of the adaptations of the Social Relations Model that were used to estimate the social integration measures. Fourth, we present a DAG (directed acyclic graph). Fifth, we describe covariates used in the models and present group demography data. Sixth, we provide plots that illustrate social integration measure change as a function of female's age. Seventh, we provide mathematical specification of Bayesian accelerated failure time model. Eighth, we report accelerated failure model results with diagnostic metrics to complement the results reported in the main text. Ninth, we present model results from accelerated failure time models with alternative specifications and priors. Tenth, we present model results from accelerated failure time models in which the previous year's social integration estimate is used to predict survival.

#### 1. PATHWAYS

Below are some possible pathways through which grooming, foraging in close proximity of others, and coalitionary aggression interactions can result in survival related benefits. These pathways are based on pathways presented by Ostner and Schülke [1] and Thompson [2]. We chose the following survival-related benefits relevant to capuchin females:

1. Reduction of risk: less wounding during coalitionary aggression
2. Reduction of risk: less predation
3. Access to food resources
4. Social status: increase in rank

Interactions in grooming, foraging, and coalitionary aggression can result in survival-related benefits as follows:

1. A female's interactions can directly result in survival related benefits
   - Feeding tolerance results in access to food resources.
   - Receiving coalitionary support results in less wounding and an increase in rank during rank-changing coalitions.
2. Female capuchins can exchange the same or different currencies with individuals in their social group
   - Grooming-giving exchanged for coalitionary support results in less wounding during coalitionary conflicts and/or an increase in rank during rank-changing coalitions [3, 4].
   - Grooming-giving exchanged for tolerance while feeding results in better access to resources [4, 5].
   - Grooming-giving in exchange for general tolerance results in less aggression and less wounding.
   - Providing coalitionary support in exchange for tolerance while feeding results in more access to food resources.

- Providing tolerance while feeding in exchange for coalitionary support results in an increase in rank during rank-changing coalitions.
- Providing coalitionary support in exchange for coalitionary support results in less wounding, and/or an increase in rank during rank-changing coalitions.

3. A female's affiliative interactions might be a proxy for overall tolerance that the female enjoys from other group members. This can manifest as more tolerance in other fitness-related interactions.

    - Feeding tolerance might be a proxy for the general tolerance of the female in other domains and might translate into less aggression and result in less wounding (inter-individual tolerance in the context of food has been linked to social bonds [4, 6] and increased levels of cooperation [7, 8].
    - Receiving grooming might be a proxy for the general tolerance of the female in other domains and translate into less aggression and less wounding.

4. A female's affiliative interactions might be a proxy for spatial centrality (the causal direction could go either way: interactions could cause centrality, or centrality might cause more interactions.)

    - Grooming (providing and receiving) is associated with spatial centrality and results in less predation.
    - Feeding tolerance (providing and receiving) is associated with spatial centrality and results in less predation.

The pathways are not mutually exclusive and the list is not exhaustive. Finding an association between one interaction type (e.g. grooming-giving) and survival is consistent with multiple pathways.

The direction, i.e. being either a provider or receiver, matters in most pathways. For example, in exchanges of coalitionary support and grooming, it matters if the female is a recipient or a giver of coalitionary support. If she is a recipient of coalitionary support then she exchanges it by providing grooming. If she is a provider of coalitionary support, then she receives grooming in exchange. In the first case, the association between grooming-giving and survival would be consistent with the pathway to survival. In the second case, the association between coalitionary support-giving and survival would be consistent with the pathway to survival. Since direction matters in the pathways, whenever possible, we separated the individual roles of giver and receiver in order to understand how social interactions translate into increased survival.

#### 2. CAUSES OF DEATH

For most females, we do not know the cause of death, because we rarely observe deaths as they occur.

Out of 62 adult female deaths that occurred between 2002-2019, we have some information about the cause for 10 females (Table S1). Five females died during a drought and their deaths might be related to the drought conditions (possibly starvation). Two females were hit by a car, one was killed by a poacher, one disappeared when she was injured, and one vanished while pregnant. For all other females, we do not have any information about the cause of their deaths, because they usually disappear while the group is not being observed.

The absence of observations of infectious disease spreading through this population suggests that infectious disease is not a prominent cause of death for capuchins.

Old age is also not a common cause of death for females. The oldest living female in this population is estimated to be 39 years old, but the maximum life span in the wild is not known, because white-faced capuchins can live up to 55 years of age in captivity [9] and no field study has lasted this long. The distribution of deaths across age categories (Table S1) suggests that females die before they reach old age .

We have no evidence for differences in mortality patterns across social groups.

**Table S1.** Causes of death. There were 132 adult females who lived in 11 social groups between 2002 and 2019. Sixty-six females died during study period (Died) and 66 were still living at the end of 2019 (Censored). The number of females that died due to the listed cause of death is provided in the parentheses.

| Age | Survived | Censored | Died | Cause of death |
|---|---|---|---|---|
| 5 | 127 | 3 | 2 | unknown (2) |
| 6 | 122 | 3 | 2 | unknown (2) |
| 7 | 113 | 7 | 2 | unknown (1), drought* (1) |
| 8 | 104 | 4 | 5 | unknown (4), drought* (1) |
| 9 | 96 | 6 | 2 | unknown (2) |
| 10 | 83 | 9 | 4 | unknown (4) |
| 11 | 76 | 3 | 4 | unknown (3), accident** (1) |
| 12 | 70 | 3 | 3 | unknown (2), accident** (1) |
| 13 | 64 | 3 | 3 | unknown (3) |
| 14 | 58 | 3 | 3 | unknown (3) |
| 15 | 48 | 5 | 5 | unknown (5) |
| 16 | 39 | 6 | 3 | unknown (3) |
| 17 | 35 | 1 | 3 | unknown (1), drought* (1), predation*** (1) |
| 18 | 31 | 3 | 1 | unknown (1) |
| 19 | 29 | 0 | 2 | unknown (2) |
| 20 | 26 | 2 | 1 | unknown (1) |
| 21 | 23 | 1 | 2 | unknown (2) |
| 22 | 21 | 1 | 1 | unknown (1) |
| 23 | 19 | 0 | 2 | unknown (2) |
| 24 | 18 | 0 | 1 | unknown (1) |
| 25 | 17 | 0 | 1 | drought* (1) |
| 26 | 13 | 2 | 2 | unknown (2) |
| 27 | 11 | 1 | 1 | unknown (1) |
| 28 | 10 | 0 | 1 | unknown (1) |
| 29 | 9 | 0 | 1 | unknown (1) |
| 30 | 9 | 0 | 0 | - |
| 31 | 9 | 0 | 0 | - |
| 32 | 9 | 0 | 0 | - |
| 33 | 8 | 0 | 1 | unknown (1) |
| 34 | 4 | 0 | 4 | unknown (3), injury**** (1) |
| 35 | 3 | 0 | 1 | drought* (1) |
| 36 | 2 | 0 | 1 | unknown (1) |
| 37 | 1 | 0 | 1 | unknown (1) |
| 38 | 1 | 0 | 0 | - |
| 39 | 0 | 0 | 1 | unknown (1) |

[]

*Death possibly due to malnutrition during a drought. **Hit by a car. ***Killed by poacher. ****Death possibly due to unknown infected injury that a monkey had for several months.

#### 3. SOCIAL INTEGRATION ESTIMATES

A central challenge to estimating sociality is the heterogeneous sampling that characterizes ethological research [10]. Because some animals and their dyadic interactions are observed more than others, there is heterogeneous uncertainty about the underlying sociality of the individuals in the population. As an alternative to point estimates of sociality, our approach was designed to obtain measures of sociality that reflect that uncertainty, which subsequently carries forward into the accelerated failure time models.

To model social integration, we adapt the multilevel Social Relations Model, or SRM [11, 12]. Originally formulated as an ANOVA model, the SRM partitions the variance of a directed dyadic outcome from actor $i$ to partner $j$ into (1) actor-level variation reflecting individual-level heterogeneity of directing or sending ties to others, (2) partner-level variation reflecting individual-level heterogeneity of receiving ties from others, and (3) dyadic variation, which reflects the extent to which the $i$ to $j$ relationship is similar, or reciprocal, to the $j$ to $i$ relationship. The multilevel parameterization of the SRM advantageously allows this variation to be modeled with random effects, or varying intercepts, which accommodates imbalanced sampling relatively easily. Originally designed for continuous responses, the SRM can be adapted for binary, count, and ordered outcomes [13–16]. Implementations of these models have become common among researchers studying cooperative behaviour in humans [17–19].

In this case, the dyadic behaviour of the capuchin monkeys is a binomial outcome, where the numerator is the number of times that the behaviour was observed and the denominator is the maximum number of times it could have been observed given the sampling of the individuals and dyads. Using a "snapshot" approach that is common to behavioural ecology [20], we aggregated the observations by calendar year. Thus, an effect for each individual and dyad was estimated on an annual basis.[1] This timeframe was chosen primarily because the accelerated failure time models employed a similar annual timeframe. Each of the behaviours (grooming, coalitional support, and foraging proximity) was estimated separately, though we note that a multivariate SRM can be used to model multiple discrete behavioural outcomes [21].

##### A. Directed behaviours

Grooming and coalitional support are directed behaviours, where the behaviour originates from individual $i$ and is directed toward individual $j$. The models assume that both individuals are members of group $k$ in year $t$. These models can be notated:

$$y_{ijkt} \sim \text{binomial}(n_{ijkt}, p_{ijkt})$$
$$\text{logit}(p_{ijkt}) = \alpha + a_{ikt} + b_{jkt} + u_{|ijkt|} + d_{ijkt}$$

where $\alpha$ represents the intercept, $a_{ikt}$ is a node-level random intercept for the sending node $i$ in group $k$ in year $t$, $b_{jkt}$ is an analogous random intercept for incoming events to $j$ in group $k$ in year $t$, $u_{|ijkt|}$ is a symmetric random intercept for dyad $ij$ in group $k$ in year $t$, and $d_{ijkt}$ is a parameter that reflects the extent to which the directed $i$ to $j$ effect is distinguished from the symmetric effect. This latter parameter also serves to reflect possible overdispersion beyond the conditional expectation for the binomial distribution [22].

The random effects for sending and recipient nodes are assumed bivariate normally distributed with zero means and a homogeneous covariance matrix:

$$\begin{pmatrix} a_{ikt} \\ b_{ikt} \end{pmatrix} \sim \text{Normal}\left\{ \begin{pmatrix} 0 \\ 0 \end{pmatrix}, \begin{pmatrix} \sigma_a^2 & \\ \sigma_{ab} & \sigma_b^2 \end{pmatrix} \right\}$$

As in other applications of the SRM [15], the correlation between the respective effects is known as the "generalized reciprocity correlation." In the present application, the modeling of this correlation helps to provide refined estimates of the node-level effects that are used as predictors in the survival analysis (i.e., the accelerated failure time models).

In practice, there is typically high correspondence between the node-level parameters and empirical calculations of out-degree and in-degree centrality [12]. That correspondence is evident

[1] Although individuals and dyads may appear in multiple group-year datasets, the random effects for each group-year were modeled separately. That is, we did not attempt to model correlations between individuals and dyads when they appear in multiple group-years.

in this study, and the primary benefit of using the Social Relations Model is to account for measurement uncertainty that stems from variation in the sampling effort of individuals.[2]

The use of symmetric dyadic effects in this analysis parallels their use by Koster et al. [23]. That is, within each dyad, the relationship from $ikt$ to $jkt$ and the relationship from $jkt$ to $ikt$ share the same index, which is distinguished by vertical pipes $|ijkt|$. The effects are assumed to be normally distributed:

$$u_{|ijkt|} \sim \text{Normal}(0, \sigma_u^2)$$

The effects for the directed dyadic effects are likewise assumed to be normally distributed:

$$d_{ijkt} \sim \text{Normal}(0, \sigma_d^2)$$

Note that binomial models lack the constant residual variance of Gaussian regression models. As an alternative, latent parameterizations of binomial models may assume that the corresponding variance is $\pi^2/3$, or 3.29 [22].

The dyadic reciprocity correlation, $\rho_{ijkt}$, can then be calculated by dividing the symmetric dyadic variance by the sum of the dyadic variances and the latent residual variance:

$$\rho_{ijkt} = \frac{\sigma_u^2}{\sigma_u^2 + \sigma_d^2 + 3.29}$$

This parameterization of $\rho_{ijkt}$ assumes that dyadic reciprocity must be positive, which we believe to be a reasonable assumption for the cooperative behaviours in this analysis.[3]

In the context of the present analysis, the inclusion of the dyadic effects mitigates the possibility that node-level estimates are disproportionately reflective of, for instance, a high volume of interactions solely within dyadic contexts. Overall, the models show very high dyadic reciprocity correlations, which further mitigates concerns that the sociality measures from the node-level random effects might be largely a by-product of asymmetric interactions between individuals.

##### B. Symmetric behaviour

Unlike grooming and coalitional support, foraging in proximity to others is regarded as an undirected, or symmetric, behaviour. For symmetric behaviour, it is not appropriate to fit a Social Relations Model because the assignment of individuals to the respective $i$ and $j$ indices is arbitrary. The random effects for these individual should have a common variance, not the respective actor-level and partner-level variances that are estimated by the SRM [12, 14, 24, 25]. [4] Individual-level variation may still be pronounced with symmetric behaviours, however, so we fit our model with individual-level random effects.

The binomial model examines the number of times that individuals $i$ and $j$ of group $k$ were observed in proximity in year $t$, denoted $y_{ijkt}$, as a proportion of the number of times that they could have been observed, $n_{ijkt}$. The model is notated as:

$$\begin{aligned}
y_{ijkt} &\sim \text{binomial}(n_{ijkt}, p_{ijkt}) \\
\text{logit}(p_{ijkt}) &= \alpha + v_{ikt} + v_{jkt} + u_{|ijkt|} \\
v_{ikt} &\sim \text{Normal}(0, \sigma_v^2) \\
v_{jkt} &\sim \text{Normal}(0, \sigma_v^2) \\
u_{|ijkt|} &\sim \text{Normal}(0, \sigma_u^2)
\end{aligned}$$

where $\alpha$ represents the intercept, $v_{ikt}$ is a node-level random intercept for the individual who has been assigned index $i$, and $v_{jkt}$ is a node-level random intercept for the individual who has been assigned index $j$. Note that the group effects are again assumed to be normally distributed with variance $\sigma_m^2$. A similar assumption applies to the individual-level effects, $v_{ikt}$ and $v_{jkt}$. Importantly, note that they share the same variance, $\sigma_v^2$. Finally, an effect for the symmetric

[2]The statistical relationship between node-level parameters and centrality measures need not be strictly monotonic [14]. That is because the model parameters are estimated jointly with other parameters in the model. In some cases, researchers might consider omitting those parameters if the goal is to obtain monotonic correspondence.

[3]Agonistic behaviours, by contrast, would potentially need to be modeled in ways that permit negative dyadic reciprocity[23].

[4]Dyadic models for symmetric data lack the nomenclature of the "Social Relations Model." Because the model we fit draws on the principles of multiple membership models [26], the approach might be described as a "Multiple Membership Relations Model."

dyad, $u_{|ijkt|}$, is included in the model to account for possible overdispersion in the variance of the binomial outcome [22].

As noted, we implemented these models in part to reflect the measurement uncertainty of heterogeneously sampled individuals. This uncertainty is evident in the individual-level estimates from the models, as seen in Figure S1. This figure, which depicts foraging proximity, represents the individuals from a single group in a single year of observation. Similar results are evident for other domains and other annualized observations of groups (not depicted here).

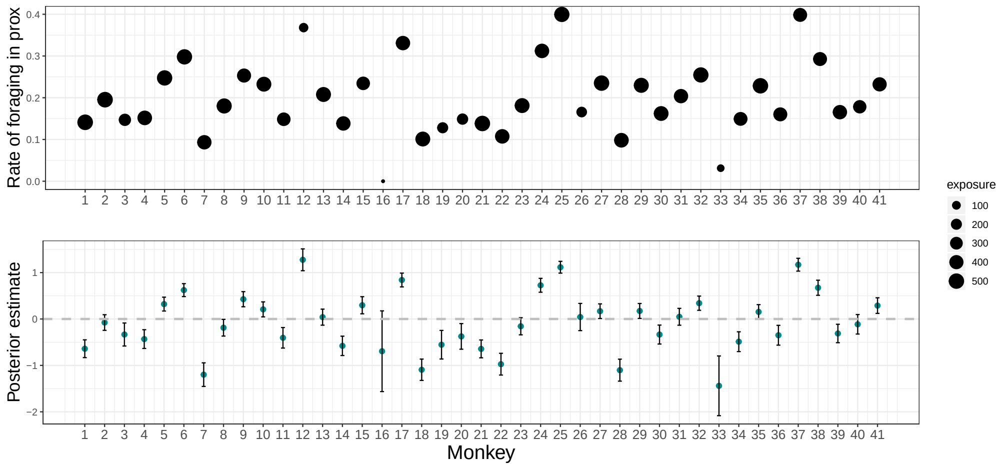


**Fig. S1.** Comparison of the empirical data to model estimates for the domain of foraging proximity for a representative group of monkeys. Points correspond to individual monkeys who were present in the group during a calendar year. The top panel shows the empirical data, represented as the rate at which individuals were observed to be foraging in proximity to others in the group. The points are sized proportional to the number of observations of that individual in that year, per the legend. The bottom panel shows the model estimates for the same monkeys, as represented by the posterior means (i.e., the points) and standard deviations (i.e., the confidence bands, which represent 1 standard deviation). Note in particular that the model estimates show greater uncertainty (larger standard deviations) as the empirical sample size decreases for a given monkey. Owing to the partial pooling, the posterior means for these individuals have also been "shrunk" toward the population average.

#### 4. DIRECTED ACYCLIC GRAPH (DAG)

Having many covariates in the model can result in "included variable bias" where predictors are not only causally influencing the outcome, but also influencing each other [27]. This can result in predictor induced statistical selection within the model and manifest itself through misleading statistical, but not causal, associations between the variables we are interested in. Our goal is to infer causal impact of social integration on survival. In addition, we considered (1) female's average dominance rank for that year, (2) her age, (3) average group size for that year, (4) the number of adult daughters co-resident with the female and (5) whether her mother was still alive.

An adult female's probability of dying in a given year is expected to increase with age [28, 29]. Females who live in larger groups might experience greater survival rates due to reduced predation risks [30] and difference in interaction networks in comparison to females in smaller social groups due to a larger number of social partners available. Higher-ranking females possibly enjoy reduced mortality in comparison to lower-ranking females due to their central position in the group [31]. A female's rank and age are likely to influence how much the female participates in social interactions. Since white-faced capuchins are a female philopatric species, and females spend their entire lives with their female relatives, we considered including the number of adult daughters co-resident that year with the female and whether her mother was still alive [28]. The presence of a mother and/or adult daughters approximates a female's close kin network benefits: females who have more extensive kin networks might experience reduced mortality [29].

To check if the above assumed relationships justify the inclusion of all of the predictor variables in the model, we drew a directed acyclical graph (DAG) using the package daggity (v.3.0) and analyzed implied functional relationships. We found that in order to test for causal impact of social integration on survival, we need to include (1) female's average dominance rank for that year, (2) age, and (3) average group size for that year.

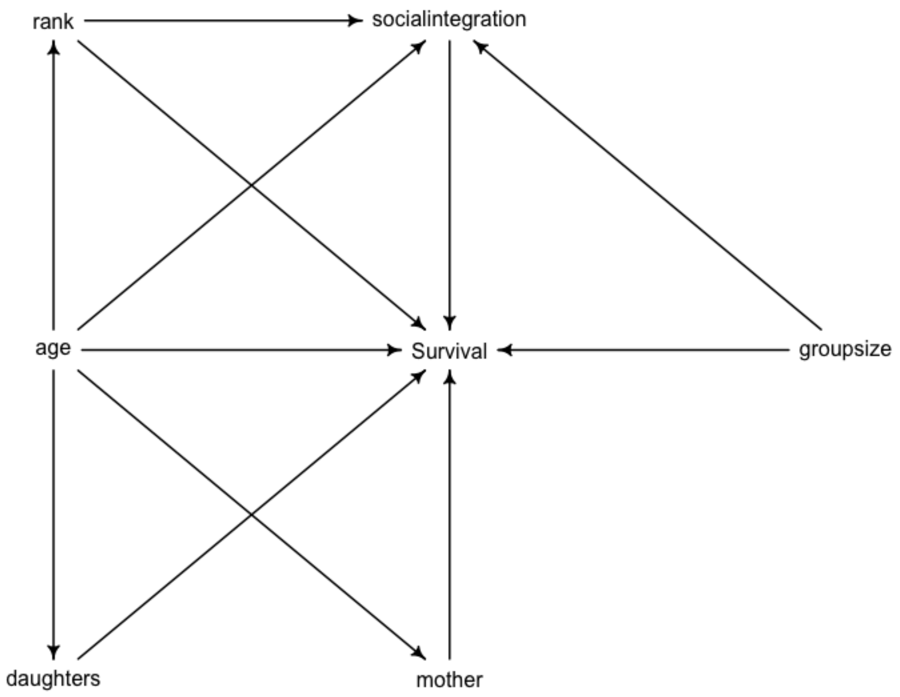


**Fig. S2.** Directed acyclic graph.

#### 5. COVARIATES

##### A. Age

Focal female ages were assigned based on demographic records of births and deaths collected from 1990 – 2019. The ages of females who were born before 1990 were estimated in part by retroactively comparing photos taken then with photos of known-aged females collected later in the study. In addition, we inferred reproductive histories via genetic maternity data. We assumed that the age of first birth for each mother was six years, and that interbirth intervals were 2 years [32, 33]. Out of 132 females in our sample, the birth year of 33 females was uncertain.

##### B. Average number of individuals in her group (group size)

This is the number of adult females, adult males, and immatures that resided in the female's group during the days when researchers spent at least six hours of observation with the group, averaged for the year. See Table S2 for group demography data.

##### C. Annual dominance index (dominance rank)

The annual dominance index represents the proportion of group members that the female dominated, on average, that year. For each observation day that the female (the focal) resided in a social group, we identified all of the other co-resident individuals (alters). To assess whether the female was dominant to an alter on a particular day, we first identified the dominance interaction immediately preceding and following the day of interest for each focal-alter dyad. Each focal-alter dyad could contribute 0, 1 or 2 data points to the daily dominance index. For each interaction, we identified that the focal individual was dominant if she was either the animal performing the supplanting, or being cowered at, or being avoided, or fled from. The daily dominance index, $DDI_i$, of a focal individual, $i$, is a sum of dominance interactions where the focal was dominant to her alters, $w_{i-a}$, divided by the total number of dominance interactions that the focal had with her alters, $s_{i-a}$:

$$DDI_i = \frac{\sum w_{i-a}}{\sum s_{i-a}}$$

Then, the average annual dominance index, $ADI_i$, is an average of daily dominance indices:

$$ADI_i = \frac{1}{n}\sum_{i=1}^{n} DDI_i$$

In some cases, either one or no dominance interactions were available for a focal-alter dyad. As a result, the individuals who did not have dominance interactions with the focal did not contribute to the calculation of the daily dominance index.

**Table S2.** Group demography. Reported quantities are a number of group members(adults and immatures) that co-resided in a social group during the days when researchers spent at least six hours of observation with the group, averaged for the year (the average number of adult females followed by the average number of adult males are in parentheses.)

| Year/ Group | AA | CE | CU | DI | FF | FL | MK | NM | RF | RR | SP |
|---|---|---|---|---|---|---|---|---|---|---|---|
| 2002 | 36 (10;10) | - | - | - | 30 (9;10) | - | - | - | - | 36 (10;11) | - |
| 2003 | 28 (7;8) | - | - | - | 30 (10;8) | 11 (4;3) | - | - | - | 37 (10;10) | - |
| 2004 | 17 (6;3) | - | - | - | 33 (11;9) | 11 (4;2) | 12 (5;2) | - | - | 30 (8;8) | - |
| 2005 | 20 (7;3) | - | - | - | 34 (11;10) | 14 (5;4) | 15 (5;3) | - | - | 25 (6;8) | 17 (5;4) |
| 2006 | 22 (8;5) | - | - | - | 37 (11;10) | 17 (5;4) | 17 (6;3) | - | - | 27 (6;8) | 13 (4;3) |
| 2007 | 22 (8;4) | - | 5 (3;2) | - | 31 (10;10) | 19 (5;6) | 16 (5;3) | - | 12 (4;2) | 27 (8;8) | - |
| 2008 | 23 (8;2) | - | 6 (3;1) | - | 18 (8;3) | 17 (4;5) | 16 (4;3) | 10 (2;3) | 16 (5;4) | 29 (8;8) | - |
| 2009 | 25 (9;2) | - | 5 (3;1) | - | 19 (9;3) | 16 (3;5) | 17 (6;3) | 14 (4;3) | 19 (7;4) | 30 (8;10) | 25 (7;7) |
| 2010 | 27 (10;3) | - | 8 (3;1) | - | 20 (9;2) | 16 (4;4) | 17 (6;5) | 15 (4;4) | 21 (7;4) | 25 (9;7) | 25 (7;6) |
| 2011 | 30 (11;3) | - | 9 (3;2) | - | 22 (9;3) | 19 (5;5) | 20 (6;5) | 14 (5;4) | 24 (10;3) | 26 (9;7) | 27 (8;7) |
| 2012 | 30 (11;3) | 7 (3;2) | 8 (1;2) | 8 (3;0) | 23 (9;3) | 19 (4;6) | 22 (6;5) | 13 (6;3) | 25 (10;2) | 28 (8;8) | 29 (8;8) |
| 2013 | 30 (9;8) | 13 (4;3) | 6 (1;1) | 11 (4;3) | 19 (7;3) | 21 (4;7) | 23 (6;6) | 15 (5;3) | 26 (11;3) | 23 (6;8) | 25 (8;7) |
| 2014 | 31 (9;9) | 14 (5;3) | - | 11 (4;2) | 17 (6;3) | 22 (4;7) | 23 (7;6) | 14 (5;3) | 28 (12;5) | 21 (6;6) | 28 (9;8) |
| 2015 | 21 (8;6) | 15 (5;6) | - | 10 (4;2) | 16 (7;4) | 23 (5;7) | 20 (6;7) | 14 (4;3) | 26 (12;5) | 21 (6;4) | 26 (10;8) |
| 2016 | 18 (8;4) | 15 (5;5) | - | 10 (4;2) | 12 (7;2) | 24 (6;6) | 18 (5;7) | 11 (3;1) | 20 (9;6) | 10 (6;1) | 26 (8;12) |
| 2017 | 22 (9;6) | 19 (5;6) | - | 10 (4;2) | 14 (7;2) | 27 (8;6) | 17 (5;5) | 13 (5;2) | 23 (9;6) | 15 (6;4) | 27 (9;9) |
| 2018 | 22 (9;6) | 18 (5;6) | - | 11 (4;2) | 16 (7;2) | 27 (9;6) | 15 (5;3) | 18 (5;4) | 25 (10;6) | 18 (6;7) | 25 (8;7) |
| 2019 | 23 (8;4) | 17 (5;5) | - | 14 (4;3) | 18 (7;3) | 30 (11;5) | 17 (5;4) | - | 26 (9;6) | 16 (4;4) | - |

#### 6. AGE AND CHANGES IN INDIVIDUAL SOCIAL INTEGRATION

In this section, we provide plots that illustrate changes in social integration measures as a function of age. Across all dataset types (all group members, adult females only, adult males only) and social integration types (*grooming giving*, *grooming receiving*, *support giving*, *support receiving*, and *foraging*) the estimates tend to be lower for older females.

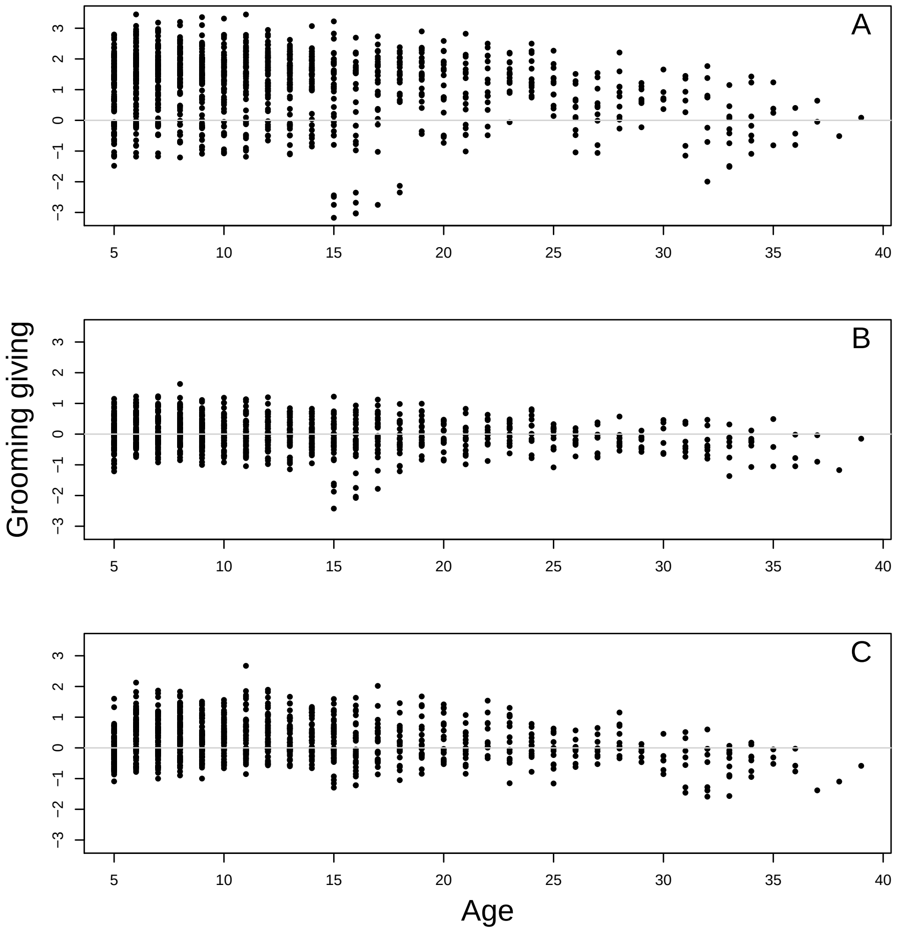


**Fig. S3.** Grooming giving posterior estimates from Social Relations Model as a function of female age: (a) all group members dataset (N=132 adult females, 1058 female years), (b) adult females only dataset (N=132 females, 985 female years), (c) adult males only dataset (N=132 females, 1002 female years).

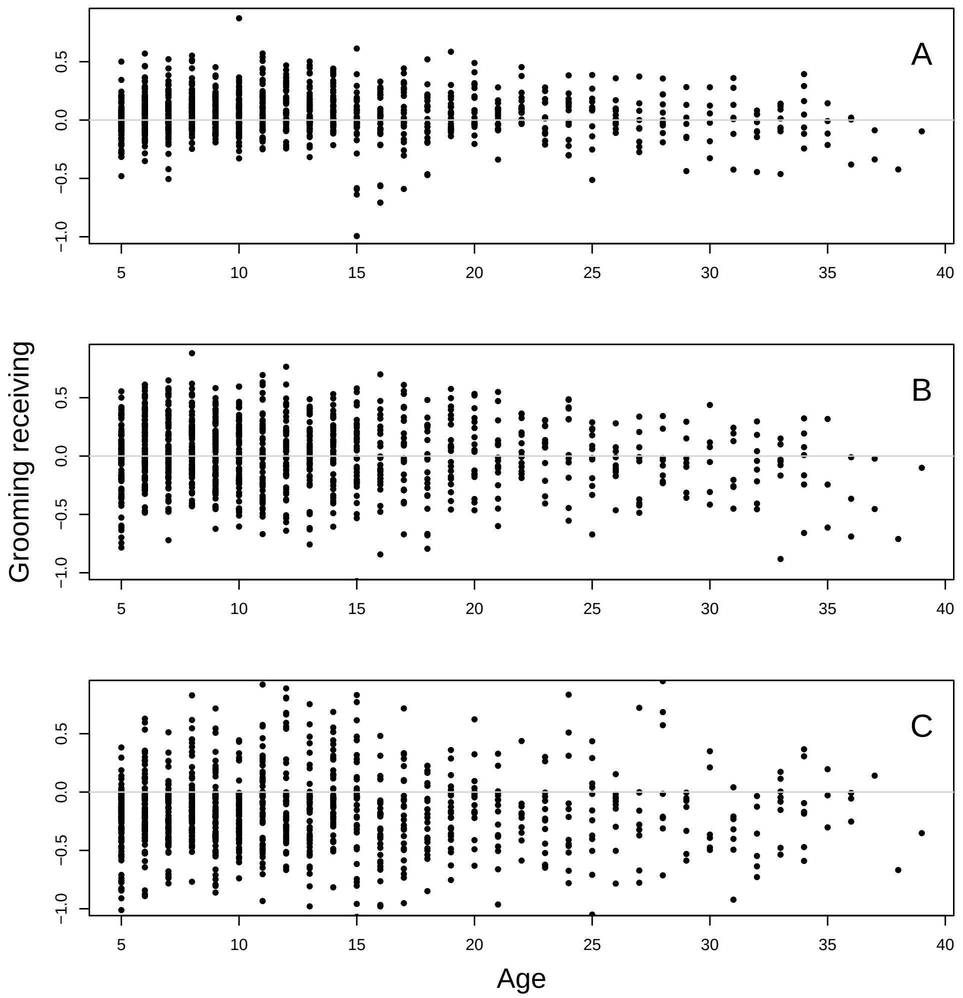


**Fig. S4.** Grooming receiving posterior estimates from Social Relations Model as a function of female age: (a) all group members dataset (N=132 adult females, 1058 female years), (b) adult females only dataset (N=132 females, 985 female years), (c) adult males only dataset (N=132 females, 1002 female years).

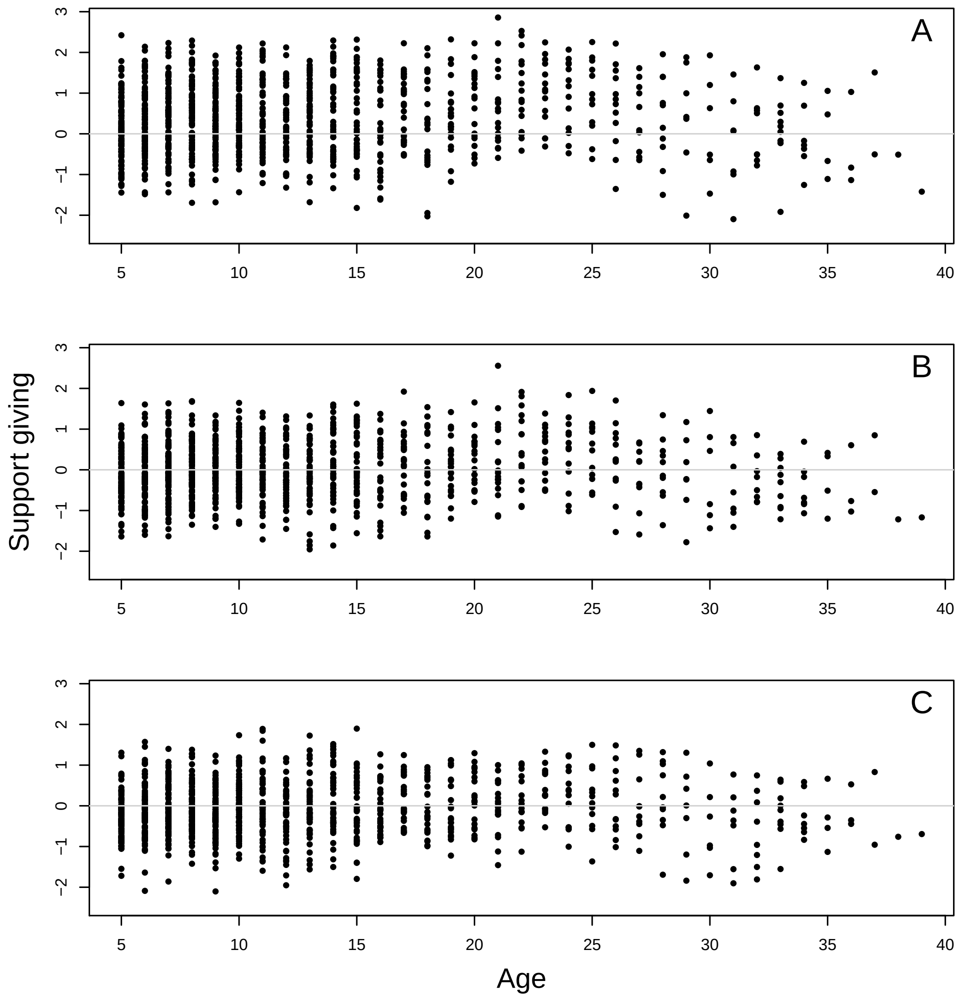


**Fig. S5.** Support giving posterior estimates from Social Relations Model as a function of female age: (a) all group members dataset (N=132 adult females, 1058 female years), (b) adult females only dataset (N=132 females, 985 female years), (c) adult males only dataset (N=132 females, 1002 female years).

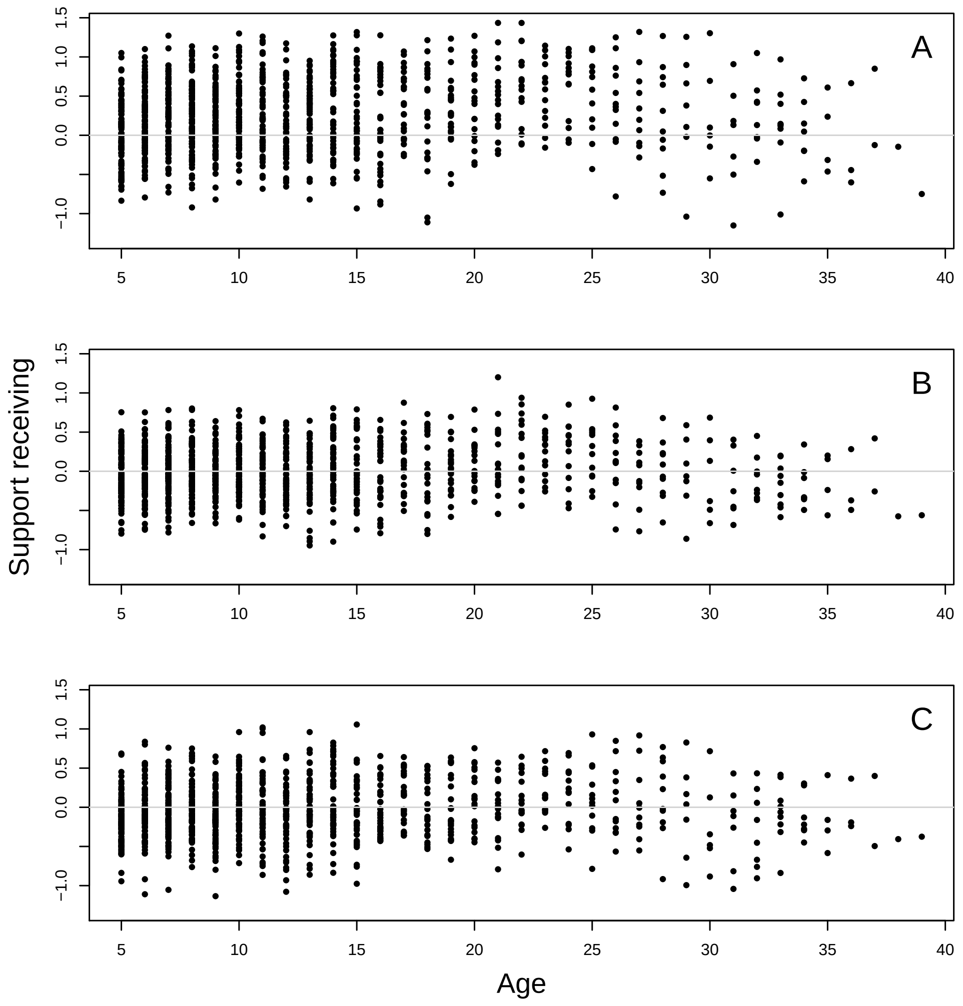


**Fig. S6.** Support receiving posterior estimates from Social Relations Model as a function of female age: (a) all group members dataset (N=132 adult females, 1058 female years), (b) adult females only dataset (N=132 females, 985 female years), (c) adult males only dataset (N=132 females, 1002 female years).

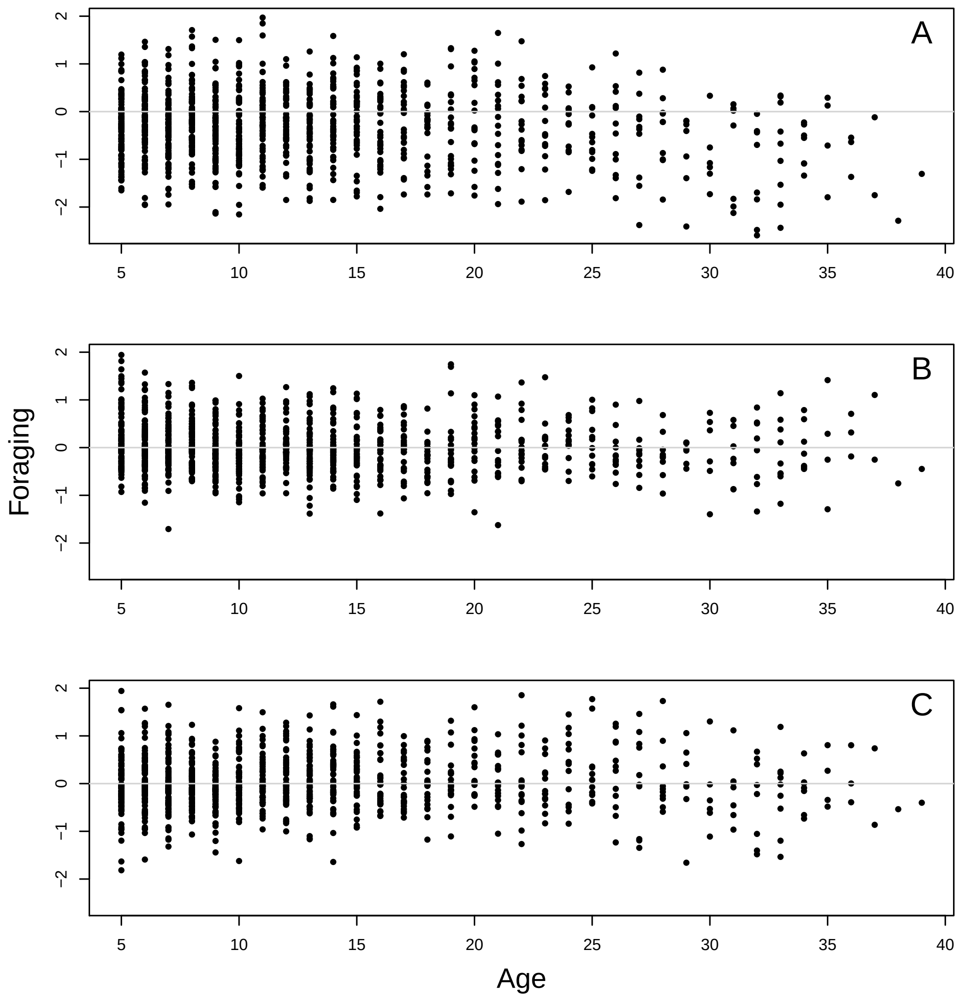


**Fig. S7.** Foraging posterior estimates from Social Relations Model as a function of female age: (a) all group members dataset (N=132 adult females, 1058 female years), (b) adult females only dataset (N=132 females, 985 female years), (c) adult males only dataset (N=132 females, 1002 female years).

#### 7. THE ACCELERATED FAILURE TIME MODEL

We specified the following model for the number of days before death, $D_i$. The probability for the number of days before death comes from the cumulative probability distribution:

$$Pr(D_i|\lambda_i) = \lambda_i exp(-\lambda_i D_i)$$

For females who did not die during the observation period, the probability of surviving $D_i$ comes from the complementary cumulative probability distribution:

$$Pr(D_i|\lambda_i) = exp(-\lambda_i D_i)$$

We model the rate of dying, $\lambda_i$, as follows:

$$\lambda_i = 1/\mu_i$$

where $\mu_i$ is the expected number of days till death

$$log(\mu_i) = \alpha + \alpha_{group}[groupid] + \alpha_{individual}[id] + b_{sociality} \cdot TrueSocialityEstimate[id] \\ +b_{rank} \cdot Rank + b_{age} \cdot Age + b_{groupsize} \cdot GroupSize$$

where $\alpha$ denotes the intercept or the base rate of number of days survived, $\alpha_{individual}[id]$ denotes individual female random effects, and $\alpha_{group}[groupid]$ denotes group random effects corresponding to the observation period. The model coefficients $b_{sociality}$ , $b_{rank}$, $b_{age}$, and $b_{groupsize}$ describe the impact of social integration, rank, age, and group size, respectively. We adopted a latent variable approach to model the social integration estimates given that the individuals' social integration estimates are not point estimates, but rather posterior distributions with means and standard deviations that reflect the measurement uncertainty.

We assumed a Normal (8, 0.5) prior for a base rate of survival, $\alpha$, which places most of the prior mass between 0 to 20 years with the mean of 8 years and a long tail allowing more extreme values. For fixed effects, we assumed Normal (0, 1) priors. For individual-level random effects, $\alpha_{individual}[id]$ and $\alpha_{group}[groupid]$, we use a Normal (0, 1) prior.

#### 8. ACCELERATED FAILURE MODEL RESULTS

Reported quantities are posterior means, standard deviation and 89% credible interval. Model diagnostic metrics provided are an estimate of the number of independent samples (n_eff) and an indicator $\hat{R}$ (Rhat4) of the convergence of the Markov chains to the target distribution.

Each table indicates which social integration measure (*grooming giving*, *grooming receiving*, *support giving*, *support receiving*, *foraging*) was used as a predictor of survival, as well as which dataset (all group members, adult females only, adult males only) and how many adult females contributed to the survival analysis.

##### A. All group members dataset

This dataset includes adult female social integration measures estimated using their interactions with all group members.

**Table S3.** Accelerated failure time model predicting female survival (N=132 females, 1058 female-years of data, 66 censored cases) as a function of *grooming giving* to all group members. Parameters in bold represent effects whose 89% credible intervals do not include zero.

| | Mean | SD | 5.5% | 94.5% | n_eff | Rhat4 |
|---|---|---|---|---|---|---|
| Intercept | **9.46** | **0.34** | **8.91** | **10.02** | 3212.95 | 1.00 |
| Grooming giving | **0.54** | **0.12** | **0.35** | **0.73** | 2976.26 | 1.00 |
| Age | **-0.07** | **0.02** | **-0.10** | **-0.04** | 3389.42 | 1.00 |
| Rank | -0.24 | 0.16 | -0.50 | 0.01 | 4724.18 | 1.00 |
| Group size | 0.15 | 0.17 | -0.11 | 0.42 | 5317.00 | 1.00 |

**Table S4.** Accelerated failure time model predicting female survival (N=132 females, 1058 female-years of data, 66 censored cases) as a function of *grooming receiving* from all group members. Parameters in bold represent effects whose 89% credible intervals do not include zero.

| | Mean | SD | 5.5% | 94.5% | n_eff | Rhat4 |
|---|---|---|---|---|---|---|
| Intercept | **9.48** | **0.33** | **8.95** | **10.01** | 2598.16 | 1.00 |
| Grooming receiving | 0.21 | 0.13 | 0.00 | 0.41 | 4739.01 | 1.00 |
| Age | **-0.09** | **0.02** | **-0.12** | **-0.06** | 2942.89 | 1.00 |
| Rank | -0.13 | 0.16 | -0.39 | 0.12 | 3566.35 | 1.00 |
| Group size | 0.12 | 0.16 | -0.15 | 0.38 | 3007.23 | 1.00 |

**Table S5.** Accelerated failure time model predicting female survival (N=132 females, 1058 female-years of data, 66 censored cases) as a function of *support giving* to all group members. Parameters in bold represent effects whose 89% credible intervals do not include zero.

| | Mean | SD | 5.5% | 94.5% | n_eff | Rhat4 |
|---|---|---|---|---|---|---|
| Intercept | **9.50** | **0.34** | **8.97** | **10.05** | 5201.03 | 1.00 |
| Support giving | 0.15 | 0.17 | -0.12 | 0.42 | 4585.91 | 1.00 |
| Age | **-0.09** | **0.02** | **-0.12** | **-0.06** | 4763.36 | 1.00 |
| Rank | -0.16 | 0.18 | -0.46 | 0.13 | 5060.26 | 1.00 |
| Group size | 0.12 | 0.16 | -0.14 | 0.38 | 6899.12 | 1.00 |

**Table S6.** Accelerated failure time model predicting female survival (N=132 females, 1058 female-years of data, 66 censored cases) as a function of *support receiving* from all group members. Parameters in bold represent effects whose 89% credible intervals do not include zero.

| | Mean | SD | 5.5% | 94.5% | n_eff | Rhat4 |
|---|---|---|---|---|---|---|
| Intercept | **9.49** | **0.34** | **8.96** | **10.03** | 3535.28 | 1.00 |
| Support receiving | 0.21 | 0.16 | -0.05 | 0.45 | 5135.50 | 1.00 |
| Age | **-0.09** | **0.02** | **-0.12** | **-0.06** | 4085.79 | 1.00 |
| Rank | -0.20 | 0.18 | -0.49 | 0.10 | 4581.18 | 1.00 |
| Group size | 0.12 | 0.16 | -0.14 | 0.38 | 4965.83 | 1.00 |

**Table S7.** Accelerated failure time model predicting female survival (N=132 females, 1058 female-years of data, 66 censored cases) as a function of *foraging* near all group members. Parameters in bold represent effects whose 89% credible intervals do not include zero.

| | Mean | SD | 5.5% | 94.5% | n_eff | Rhat4 |
|---|---|---|---|---|---|---|
| Intercept | **9.47** | **0.34** | **8.92** | **10.02** | 2565.89 | 1.00 |
| Foraging | 0.28 | 0.18 | -0.01 | 0.58 | 2219.40 | 1.00 |
| Age | **-0.09** | **0.02** | **-0.12** | **-0.06** | 2751.32 | 1.00 |
| Rank | -0.18 | 0.17 | -0.45 | 0.09 | 3364.26 | 1.00 |
| Group size | 0.22 | 0.18 | -0.07 | 0.51 | 3622.56 | 1.00 |

##### B. Adult females only dataset

This dataset includes adult female social integration measures estimated using their interactions with adult females only.

**Table S8.** Accelerated failure time model predicting female survival (N=132 females, 985 female-years of data, 77 censored cases) as a function of *grooming giving* to adult females only. Parameters in bold represent effects whose 89% credible intervals do not include zero.

| | Mean | SD | 5.5% | 94.5% | n_eff | Rhat4 |
|---|---|---|---|---|---|---|
| Intercept | **9.41** | **0.34** | **8.88** | **9.96** | 3870.66 | 1.00 |
| Grooming giving | **0.30** | **0.15** | **0.07** | **0.54** | 4658.88 | 1.00 |
| Age | **-0.07** | **0.02** | **-0.10** | **-0.04** | 3672.11 | 1.00 |
| Rank | -0.14 | 0.17 | -0.41 | 0.13 | 4693.99 | 1.00 |
| Group size | 0.09 | 0.18 | -0.18 | 0.38 | 4809.67 | 1.00 |

**Table S9.** Accelerated failure time model predicting female survival (N=132 females, 985 female-years of data, 77 censored cases) as a function of *grooming receiving* from adult females only. Parameters in bold represent effects whose 89% credible intervals do not include zero.

| | Mean | SD | 5.5% | 94.5% | n_eff | Rhat4 |
|---|---|---|---|---|---|---|
| Intercept | **9.43** | **0.34** | **8.89** | **9.99** | 2587.80 | 1.00 |
| Grooming receiving | **0.29** | **0.14** | **0.07** | **0.51** | 4377.72 | 1.00 |
| Age | **-0.07** | **0.02** | **-0.10** | **-0.05** | 2594.34 | 1.00 |
| Rank | -0.15 | 0.17 | -0.42 | 0.12 | 3555.30 | 1.00 |
| Group size | 0.09 | 0.17 | -0.18 | 0.38 | 4449.75 | 1.00 |

**Table S10.** Accelerated failure time model predicting female survival (N=132 females, 985 female-years of data, 77 censored cases) as a function of *support giving* to adult females only. Parameters in bold represent effects whose 89% credible intervals do not include zero.

| | Mean | SD | 5.5% | 94.5% | n_eff | Rhat4 |
|---|---|---|---|---|---|---|
| Intercept | **9.46** | **0.34** | **8.94** | **10.02** | 3461.09 | 1.00 |
| Support giving | 0.25 | 0.17 | -0.03 | 0.53 | 3558.00 | 1.00 |
| Age | **-0.08** | **0.02** | **-0.11** | **-0.05** | 2814.80 | 1.00 |
| Rank | -0.19 | 0.19 | -0.48 | 0.11 | 3860.90 | 1.00 |
| Group size | 0.08 | 0.18 | -0.20 | 0.36 | 5074.32 | 1.00 |

**Table S11.** Accelerated failure time model predicting female survival (N=132 females, 985 female-years of data, 77 censored cases) as a function of *support receiving* from adult females only. Parameters in bold represent effects whose 89% credible intervals do not include zero.

| | Mean | SD | 5.5% | 94.5% | n_eff | Rhat4 |
|---|---|---|---|---|---|---|
| Intercept | **9.45** | **0.35** | **8.89** | **9.99** | 3269.30 | 1.00 |
| Support receiving | 0.23 | 0.17 | -0.04 | 0.51 | 4907.51 | 1.00 |
| Age | **-0.08** | **0.02** | **-0.11** | **-0.05** | 3568.05 | 1.00 |
| Rank | -0.18 | 0.19 | -0.47 | 0.13 | 3747.50 | 1.00 |
| Group size | 0.08 | 0.17 | -0.19 | 0.35 | 4297.86 | 1.00 |

**Table S12.** Accelerated failure time model predicting female survival (N=132 females, 985 female-years of data, 77 censored cases) as a function of *foraging* near adult females only. Parameters in bold represent effects whose 89% credible intervals do not include zero.

| | Mean | SD | 5.5% | 94.5% | n_eff | Rhat4 |
|---|---|---|---|---|---|---|
| Intercept | **9.48** | **0.34** | **8.94** | **10.01** | 3600.79 | 1.00 |
| Foraging | **0.32** | **0.16** | **0.06** | **0.58** | 2679.56 | 1.00 |
| Age | **-0.08** | **0.02** | **-0.11** | **-0.05** | 3576.73 | 1.00 |
| Rank | -0.15 | 0.18 | -0.43 | 0.12 | 4130.90 | 1.00 |
| Group size | 0.13 | 0.18 | -0.16 | 0.42 | 5114.30 | 1.00 |

##### C. Adult males only dataset

This dataset includes adult female social integration measures estimated using their interactions with adult males only.

**Table S13.** Accelerated failure time model predicting female survival (N=29 females, 1002 female-years of data, 74 censored cases) as a function of *grooming giving* near adult males only. Parameters in bold represent effects whose 89% credible intervals do not include zero.

| | Mean | SD | 5.5% | 94.5% | n_eff | Rhat4 |
|---|---|---|---|---|---|---|
| Intercept | **9.46** | **0.34** | **8.92** | **10.01** | 4529.20 | 1.00 |
| Grooming giving | 0.20 | 0.15 | -0.04 | 0.45 | 3950.04 | 1.00 |
| Age | **-0.08** | **0.02** | **-0.11** | **-0.05** | 4764.72 | 1.00 |
| Rank | -0.05 | 0.16 | -0.31 | 0.21 | 5488.66 | 1.00 |
| Group size | 0.10 | 0.17 | -0.17 | 0.37 | 6817.84 | 1.00 |

**Table S14.** Accelerated failure time model predicting female survival (N=29 females, 1002 female-years of data, 74 censored cases) as a function of *grooming receiving* near adult males only. Parameters in bold represent effects whose 89% credible intervals do not include zero.

| | Mean | SD | 5.5% | 94.5% | n_eff | Rhat4 |
|---|---|---|---|---|---|---|
| Intercept | **9.52** | **0.34** | **8.96** | **10.06** | 2784.14 | 1.00 |
| Grooming receiving | -0.10 | 0.13 | -0.31 | 0.10 | 3430.95 | 1.00 |
| Age | **-0.09** | **0.02** | **-0.12** | **-0.06** | 3058.89 | 1.00 |
| Rank | 0.02 | 0.16 | -0.24 | 0.28 | 4402.97 | 1.00 |
| Group size | 0.07 | 0.17 | -0.21 | 0.34 | 3942.03 | 1.00 |

**Table S15.** Accelerated failure time model predicting female survival (N=29 females, 1002 female-years of data, 74 censored cases) as a function of *support giving* near adult males only. Parameters in bold represent effects whose 89% credible intervals do not include zero.

| | Mean | SD | 5.5% | 94.5% | n_eff | Rhat4 |
|---|---|---|---|---|---|---|
| Intercept | **9.52** | **0.34** | **8.98** | **10.07** | 2854.34 | 1.00 |
| Support giving | -0.14 | 0.16 | -0.40 | 0.12 | 2954.79 | 1.00 |
| Age | **-0.09** | **0.02** | **-0.12** | **-0.06** | 3467.09 | 1.00 |
| Rank | 0.07 | 0.18 | -0.20 | 0.35 | 3691.44 | 1.00 |
| Group size | 0.05 | 0.17 | -0.23 | 0.34 | 3553.75 | 1.00 |

**Table S16.** Accelerated failure time model predicting female survival (N=29 females, 1002 female-years of data, 74 censored cases) as a function of *support receiving* near adult males only. Parameters in bold represent effects whose 89% credible intervals do not include zero.

| | Mean | SD | 5.5% | 94.5% | n_eff | Rhat4 |
|---|---|---|---|---|---|---|
| Intercept | **9.53** | **0.34** | **8.98** | **10.07** | 3519.99 | 1.00 |
| Support receiving | -0.12 | 0.15 | -0.38 | 0.12 | 4460.92 | 1.00 |
| Age | **-0.09** | **0.02** | **-0.12** | **-0.06** | 3620.38 | 1.00 |
| Rank | 0.07 | 0.17 | -0.21 | 0.34 | 4263.88 | 1.00 |
| Group size | 0.06 | 0.18 | -0.22 | 0.35 | 4731.93 | 1.00 |

**Table S17.** Accelerated failure time model predicting female survival (N=29 females, 1002 female-years of data, 74 censored cases) as a function of *foraging* near adult males only. Parameters in bold represent effects whose 89% credible intervals do not include zero.

| | Mean | SD | 5.5% | 94.5% | n_eff | Rhat4 |
|---|---|---|---|---|---|---|
| Intercept | **9.51** | **0.35** | **8.95** | **10.07** | 2225.40 | 1.00 |
| Foraging | 0.08 | 0.16 | -0.17 | 0.34 | 2184.88 | 1.00 |
| Age | **-0.09** | **0.02** | **-0.11** | **-0.06** | 2399.00 | 1.00 |
| Rank | -0.02 | 0.17 | -0.30 | 0.25 | 2962.19 | 1.00 |
| Group size | 0.10 | 0.17 | -0.17 | 0.38 | 3099.34 | 1.00 |

#### 9. ALTERNATIVE SPECIFICATIONS OF THE ACCELERATED FAILURE TIME MODELS

In this section, we present model results when we alter the parameterizations of the accelerated failure time models.

We compared three types of models for the dataset containing adult female interactions of all group members: one with no random effects and two with individual-level random effect priors assumed as Normal (0, 0.2) or Normal (0, 1). All three model types resulted in similar estimates; therefore here we are reporting the results with individual-level random effect prior Normal (0, 1) in the main text, while the results of the other two types of models are reported here (Table S18 and Table S19).

**Table S18.** Models without random effects. Each column represents the accelerated failure time model where social integration measure is either *grooming giving*, or *grooming receiving*, or *support giving*, or *support receiving*, or *foraging*. Estimates of fixed effects of each of the accelerated failure time models: posterior means and 89% HPDI.

| Parameters | Grooming giving | Grooming receiving | Support giving | Support receiving | Foraging |
|---|---|---|---|---|---|
| Intercept | 8.96 (8.72; 9.21) | 8.91 (8.67; 9.19) | 8.84 (8.62; 9.07) | 8.85 (8.63; 9.07) | 8.85 (8.62; 9.09) |
| Social integration | 0.45 (0.27; 0.61) | 0.20 (-0.01; 0.41) | 0.10 (-0.18; 0.37) | 0.18 (-0.07; 0.42) | 0.09 (-0.20; 0.39) |
| Rank | -0.16 (-0.37; 0.03) | -0.10 (-0.30; 0.10) | -0.09 (-0.34; 0.18) | -0.14 (-0.39; 0.11) | -0.06 (-0.30; 0.18) |
| Age | -0.45 (-0.61; -0.28) | -0.52 (-0.68; -0.35) | -0.54 (-0.70; -0.38) | -0.54 (-0.70; -0.38) | -0.53 (-0.71; -0.36) |
| Group size | 0.20 (-0.01; 0.41) | 0.19 (-0.01; 0.39) | 0.19 (-0.01; 0.39) | 0.20 (-0.02; 0.41) | 0.22 (-0.03; 0.47) |

**Table S19.** Models with random effects prior Normal(0, 0.2). Each column represents the accelerated failure time model where social integration measure is either *grooming giving*, or *grooming receiving*, or *support giving*, or *support receiving*, or *foraging*. Estimates of fixed effects of each of the accelerated failure time models: posterior means and 89% HPDI.

| Parameters | Grooming giving | Grooming receiving | Support giving | Support receiving | Foraging |
|---|---|---|---|---|---|
| Intercept | 9.14 (8.84; 9.44) | 9.08 (8.77; 9.41) | 8.99 (8.71; 9.28) | 9.01 (8.72; 9.30) | 9.00 (8.72; 9.30) |
| Social integration | 0.50 (0.32; 0.69) | 0.21 (-0.02; 0.44) | 0.10 (-0.19; 0.38) | 0.21 (-0.06; 0.46) | 0.14 (-0.16; 0.43) |
| Rank | -0.20 (-0.46; 0.05) | -0.10 (-0.37; 0.16) | -0.10 (-0.41; 0.20) | -0.17 (-0.48; 0.13) | -0.10 (-0.37; 0.17) |
| Age | -0.67 (-0.89; -0.44) | -0.79 (-1.01; -0.56) | -0.81 (-1.02; -0.59) | -0.81 (-1.03; -0.59) | -0.79 (-1.02; -0.57) |
| Group size | 0.16 (-0.02; 0.34) | 0.15 (-0.03; 0.34) | 0.15 (-0.04; 0.34) | 0.15 (-0.04; 0.34) | 0.18 (-0.02; 0.39) |

#### 10. ASSOCIATION BETWEEN PREVIOUS YEAR'S SOCIAL INTEGRATION ESTIMATE AND SURVIVAL

In this section, we present the model results when we model survival in year *t* as a function of the female's social integration measure in the previous year, *t-1*. The sample sizes of the datasets for these analyses are smaller than the datasets reported in the main text because in some cases behavioral data from the previous year were not available.

Table S20 shows the results for the dataset containing female interactions with all group members. For this analysis, all groups contributed at least 6 calendar years of data (mean = 12.3, SD = 4.3, range: 6 - 17). The subjects of this analysis were 125 females (899 female years), of whom 59 died during the observation period.

Table S21 shows the results for the dataset containing female interactions with females only. For this analysis, all groups contributed at least 6 calendar years of data (mean = 11.5, SD = 4.4, range: 6 - 17). The subjects of this analysis were 125 females (835 female years), of whom 55 died during the observation period.

Table S22 shows the results for the dataset containing female interactions with males only. For this analysis, all groups contributed at least 6 calendar years of data (mean = 12.1, SD = 4.2, range: 6 - 17). The subjects of this analysis were 125 females (882 female years), of whom 59 died during the observation period.

Overall, the inferences from this analysis are comparable to the inferences from the models reported in the main text.

**Table S20.** Models with female's previous year's social integration measured through interactions with all group members. Each column represents the accelerated failure time model where social integration measure is either *grooming giving*, or *grooming receiving*, or *support giving*, or *support receiving*, or *foraging*. The reported quantities are posterior means (89% credible intervals in parentheses).

| Parameters | Grooming giving | Grooming receiving | Support giving | Support receiving | Foraging |
|---|---|---|---|---|---|
| Intercept | 8.72 (8.22; 9.20) | 8.72 (8.24; 9.20) | 8.68 (8.19; 9.17) | 8.68 (8.20; 9.16) | 8.68 (8.21; 9.15) |
| Social integration | 0.28 (0.06; 0.51) | 0.13 (-0.11; 0.38) | 0.04 (-0.24; 0.32) | 0.06 (-0.21; 0.32) | 0.15 (-0.13; 0.44) |
| Age | -0.81 (-1.06; -0.56) | -0.88 (-1.12; -0.64) | -0.90 (-1.13; -0.67) | -0.90 (-1.14; -0.67) | -0.88 (-1.12; -0.64) |
| Rank | -0.13 (-0.41; 0.15) | -0.07 (-0.35; 0.21) | -0.05 (-0.38; 0.28) | -0.07 (-0.38; 0.24) | -0.09 (-0.38; 0.19) |
| Group size | 0.18 (-0.10; 0.45) | 0.16 (-0.11; 0.44) | 0.16 (-0.11; 0.43) | 0.16 (-0.12; 0.45) | 0.19 (-0.09; 0.48) |

**Table S21.** Models with female's previous year's social integration measured through interactions with females only. Each column represents the accelerated failure time model where social integration measure is either *grooming giving*, or *grooming receiving*, or *support giving*, or *support receiving*, or *foraging*. The reported quantities are posterior means (89% credible intervals in parentheses).

| Parameters | Grooming giving | Grooming receiving | Support giving | Support receiving | Foraging |
|---|---|---|---|---|---|
| Intercept | 8.69 (8.13; 9.27) | 8.72 (8.13; 9.32) | 8.68 (8.10; 9.27) | 8.67 (8.09; 9.25) | 8.70 (8.09; 9.29) |
| Social integration | 0.21 (-0.12; 0.53) | 0.22 (-0.10; 0.53) | 0.16 (-0.20; 0.51) | 0.15 (-0.20; 0.48) | 0.10 (-0.22; 0.41) |
| Age | -0.83 (-1.12; -0.53) | -0.85 (-1.15; -0.55) | -0.88 (-1.17; -0.58) | -0.88 (-1.18; -0.59) | -0.88 (-1.17; -0.58) |
| Rank | -0.05 (-0.41; 0.29) | -0.06 (-0.42; 0.28) | -0.09 (-0.49; 0.32) | -0.08 (-0.48; 0.32) | -0.02 (-0.38; 0.33) |
| Group size | 0.11 (-0.23; 0.45) | 0.12 (-0.23; 0.46) | 0.10 (-0.26; 0.45) | 0.10 (-0.25; 0.46) | 0.10 (-0.26; 0.46) |

**Table S22.** Models with female's previous year's social integration measured through interactions with males only. Each column represents the accelerated failure time model where social integration measure is either *grooming giving*, or *grooming receiving*, or *support giving*, or *support receiving*, or *foraging*. The reported quantities are posterior means (89% credible intervals in parentheses).

| Parameters | Grooming giving | Grooming receiving | Support giving | Support receiving | Foraging |
|---|---|---|---|---|---|
| Intercept | 8.68 (8.11; 9.27) | 8.72 (8.14; 9.31) | 8.68 (8.09; 9.25) | 8.67 (8.10; 9.26) | 8.67 (8.09; 9.25) |
| Social integration | 0.01 (-0.29; 0.30) | 0.16 (-0.10; 0.41) | -0.04 (-0.37; 0.27) | -0.03 (-0.36; 0.28) | 0.09 (-0.21; 0.42) |
| Age | -0.90 (-1.21; -0.57) | -0.89 (-1.19; -0.60) | -0.90 (-1.20; -0.61) | -0.90 (-1.20; -0.61) | -0.9 (-1.20; -0.6) |
| Rank | -0.04 (-0.39; 0.30) | -0.05 (-0.37; 0.27) | -0.02 (-0.39; 0.34) | -0.03 (-0.38; 0.31) | -0.08 (-0.45; 0.28) |
| Group size | 0.11 (-0.23; 0.46) | 0.13 (-0.21; 0.47) | 0.10 (-0.25; 0.45) | 0.10 (-0.25; 0.45) | 0.12 (-0.22; 0.48) |
